# Zn^2+^ is Essential for Ca^2+^ Oscillations in Mouse Eggs

**DOI:** 10.1101/2023.04.13.536745

**Authors:** Hiroki Akizawa, Emily Lopes, Rafael A. Fissore

## Abstract

Changes in the intracellular concentration of free calcium (Ca^2+^) underpin egg activation and initiation of development in animals and plants. In mammals, the Ca^2+^ release is periodical, known as Ca^2+^ oscillations, and mediated by the type 1 inositol 1,4,5-trisphosphate receptor (IP_3_R1). Another divalent cation, zinc (Zn^2+^), increases exponentially during oocyte maturation and is vital for meiotic transitions, arrests, and polyspermy prevention. It is unknown if these pivotal cations interplay during fertilization. Here, using mouse eggs, we showed that basal concentrations of labile Zn^2+^ are indispensable for sperm-initiated Ca^2+^ oscillations because Zn^2+^-deficient conditions induced by cell-permeable chelators abrogated Ca^2+^ responses evoked by fertilization and other physiological and pharmacological agonists. We also found that chemically- or genetically generated eggs with lower levels of labile Zn^2+^ displayed reduced IP_3_R1 sensitivity and diminished ER Ca^2+^ leak despite the stable content of the stores and IP_3_R1 mass. Resupplying Zn^2+^ restarted Ca^2+^ oscillations, but excessive Zn^2+^ prevented and terminated them, hindering IP_3_R1 responsiveness. The findings suggest that a window of Zn^2+^ concentrations is required for Ca^2+^ responses and IP_3_R1 function in eggs, ensuring optimal response to fertilization and egg activation.

## Introduction

Vertebrate eggs are arrested at the metaphase stage of the second meiosis (MII) when ovulated because they have an active Cdk1/cyclin B complex and inactive APC/C^Cdc20^ (Heim et al., 2018). Release from MII initiates egg activation, the first hallmark of embryonic development (Ducibella et al., 2002; Schultz and Kopf, 1995). The universal signal of egg activation is an increase in the intracellular concentration of calcium (Ca^2+^) (Ridgway et al., 1977; Stricker, 1999). Ca^2+^ release causes the inactivation of the APC/C inhibitor Emi2, which enhances cyclin B degradation and induces meiotic exit (Lorca et al., 1993; Shoji et al., 2006; Suzuki et al., 2010a). In mammals, the stereotypical fertilization Ca^2+^ signal, oscillations, consists of transient but periodical Ca^2+^ increases that promote progression into interphase (Deguchi et al., 2000; Miyazaki et al., 1986). The sperm-borne Phospholipase C zeta1 (PLCζ) persistently stimulates the production of inositol 1,4,5-trisphosphate (IP_3_) (Matsu-ura et al., 2019; Saunders et al., 2002; Wu et al., 2001) that binds its cognate receptor in the endoplasmic reticulum (ER), IP_3_R1 and causes Ca^2+^ release from the egg’s main Ca^2+^ reservoir (Wakai et al., 2019). The intake of extracellular Ca^2+^ via plasma membrane channels and transporters ensures the persistence of the oscillations (Miao et al., 2012; Stein et al., 2020; Wakai et al., 2019, 2013).

Before fertilization, maturing oocytes undergo cellular and biochemical modifications (see for review (Ajduk et al., 2008)). The nucleus of immature oocytes, known as the germinal vesicle (GV), undergoes the breakdown of its envelope marking the onset of maturation and setting in motion a series of cellular events that culminate with the release of the first polar body, the correct ploidy for fertilization, and re-arrest at MII (Eppig, 1996). Other organelles are also reorganized, such as cortical granules migrate to the cortex for exocytosis and polyspermy block, mitochondria undergo repositioning, and the cytoplasm’s redox state becomes progressively reduced to promote the exchange of the sperm’s protamine load (Liu, 2011; Perreault et al., 1988; Wakai et al., 2014). Wide-ranging adaptations also occur in the Ca^2+^ release machinery to produce timely and protracted Ca^2+^ oscillations following sperm entry (Fujiwara et al., 1993; Lawrence et al., 1998), including the increase in the content of the Ca^2+^ stores, ER reorganization with cortical cluster formation, and increased IP_3_R1 sensitivity (Lee et al., 2006; Wakai et al., 2012). The total intracellular levels of zinc (Zn^2+^) also remarkably increase during maturation, amounting to a 50% rise, which is necessary for oocytes to proceed to the telophase I of meiosis and beyond (Kim et al., 2010). Remarkably, after fertilization, Zn^2+^ levels need to decrease, as Emi2 is a Zn^2+^-associated molecule, and high Zn^2^ levels prevent MII exit (Bernhardt et al., 2012; Shoji et al., 2014; Suzuki et al., 2010b). Following the initiation of Ca^2+^ oscillations, approximately 10 to 20% of the Zn^2+^ accrued during maturation is ejected during the Zn^2+^ sparks, a conserved event in vertebrates and invertebrate species (Converse and Thomas, 2020; Kim et al., 2011; Mendoza et al., 2022; Que et al., 2019; Seeler et al., 2021; Tokuhiro and Dean, 2018; Wozniak et al., 2020; Zhang et al., 2016). The use of Zn^2+^ chelators such as N,N,N,N-tetrakis (2-pyridinylmethyl)-1,2-ethylenediamine (TPEN) to create Zn^2+^-deficient conditions buttressed the importance of Zn^2+^ during meiotic transitions (Kim et al., 2010; Suzuki et al., 2010b). However, whether the analogous dynamics of Ca^2+^ and Zn^2+^ during maturation imply crosstalk and Zn^2+^ levels modulate Ca^2+^ release during fertilization is unknown.

IP_3_Rs are the most abundant intracellular Ca^2+^ release channel in non-muscle cells (Berridge, 2016). They form a channel by assembling into tetramers with each subunit of ∼270kDa MW (Taylor and Tovey, 2010). Mammalian eggs express the type I IP_3_R, the most widespread isoform (Fissore et al., 1999; Parrington et al., 1998). IP_3_R1 is essential for egg activation because its inhibition precludes Ca^2+^ oscillations (Miyazaki and Ito, 2006; Miyazaki et al., 1992; Xu et al., 2003). Myriad and occasionally cell-specific factors influence Ca^2+^ release through the IP_3_R1 (Taylor and Tovey, 2010). For example, following fertilization, IP_3_R1 undergoes ligand-induced degradation caused by the sperm-initiated long-lasting production of IP_3_ that effectively reduces the IP_3_R1 mass (Brind et al., 2000; Jellerette et al., 2000). Another regulatory mechanism is Ca^2+^, a universal cofactor, which biphasically regulates IP_3_Rs’ channel opening (Iino, 1990; Jean and Klee, 1986), congruent with several Ca^2+^ and calmodulin binding sites on the channel’s sequence (Sienaert et al., 1997; Sipma et al., 1999). Notably, Zn^2+^ may also participate in IP_3_R1 regulation. Recent studies using electron cryomicroscopy (cryoEM), a technique that allows peering into the structure of IP_3_R1 with a near-atomic resolution, have revealed that a helical linker (LNK) domain near the C-terminus mediates the coupling between the N- and C-terminal ends necessary for channel opening (Fan et al., 2015). The LNK domain contains a putative Zinc-finger motif proposed to be vital for IP_3_R1 function (Fan et al., 2015; Paknejad and Hite, 2018). Therefore, the exponential increase in Zn^2+^ levels in maturing oocytes, besides its essential role in meiosis progression, may optimize the IP_3_R1 function, revealing hitherto unknown cooperation between these cations during fertilization.

Here, we examined whether crosstalk between Ca^2+^ and Zn^2+^ is required to initiate and sustain Ca^2+^ oscillations and maintain Ca^2+^ store content in MII eggs. We found that Zn^2+^-deficient conditions inhibited Ca^2+^ release and oscillations without reducing Ca^2+^ stores, IP_3_ production, IP_3_R1 expression, or altering the viability of eggs or zygotes. We show instead that Zn^2+^ deficiency impaired IP_3_R1 function and lessened the receptor’s ability to gate Ca^2+^ release out of the ER. Remarkably, resupplying Zn^2+^ re-established the oscillations interrupted by low Zn^2+^, although persistent increases in intracellular Zn^2+^ were harmful, disrupting the Ca^2+^ responses and preventing egg activation. Together, the results show that besides contributing to oocyte maturation, Zn^2+^ has a central function in Ca^2+^ homeostasis such that optimal Zn^2+^ concentrations ensure IP_3_R1 function and the Ca^2+^ oscillations required for initiating embryo development.

## Results

### TPEN dose-dependently lowers intracellular Zn_2+_ and inhibits sperm-initiated Ca_2+_ oscillations

TPEN is a cell-permeable, non-specific chelator with a high affinity for transition metals widely used to study their function in cell physiology (Arslan et al., 1985; Lo et al., 2020). Mouse oocytes and eggs have exceedingly high intracellular concentrations of Zn^2+^ (Kim et al., 2011, 2010), and the TPEN-induced defects in the progression of meiosis have been ascribed to its chelation (Bernhardt et al., 2011; Kim et al., 2010). In support of this view, the Zn^2+^ levels of cells showed acute reduction after TPEN addition, as reported by indicators such as FluoZin-3 (Arslan et al., 1985; Gee et al., 2002; Suzuki et al., 2010b). Studies in mouse eggs also showed that the addition of low µM (40-100) concentrations of TPEN disrupted Ca^2+^ oscillations initiated by fertilization or SrCl_2_ (Lawrence et al., 1998; Suzuki et al., 2010b), but the mechanism(s) and target(s) of the inhibition remained unknown. To gain insight into this phenomenon, we first performed dose-titration studies to determine the effectiveness of TPEN in lowering Zn^2+^ in eggs. The addition of 2.5 µM TPEN protractedly reduced Zn^2+^ levels, whereas 5 and 10 µM TPEN acutely and persistently reduced FluoZin-3 fluorescence (**Fig. 1A)**. These concentrations of TPEN are higher than the reported free Zn^2+^ concentrations in cells, but within range of those of found in typical culture conditions (Lo et al., 2020; Qin et al., 2011). We next determined the concentrations of TPEN required to abrogate fertilization-initiated oscillations. Following intracytoplasmic sperm injection (ICSI), we monitored Ca^2+^ responses while increasing TPEN concentrations. As shown in **Fig. 1B**, 5 and 10 µM TPEN effectively blocked ICSI-induced Ca^2+^ oscillations in over half of the treated cells, and the remaining eggs, after a prolonged interval, resumed lower-frequency rises (**Fig. 1B**-center panels). Finally, 50 µM or greater concentrations of TPEN permanently blocked these oscillations (**Fig. 1B**-right panel). It is noteworthy that at the time of addition, TPEN concentrations of 5 µM or above induce a sharp drop in basal Fura-2 F340/ F380 ratios, consistent with Fura-2’s high affinity for Zn^2+^ (Snitsarev et al., 1996). We next used membrane-permeable and -impermeable chelators to assess whether TPEN inhibited Ca^2+^ oscillations by chelating Zn^2+^ from intracellular or extracellular compartments. The addition of the high-affinity but cell-impermeable Zn^2+^ chelators DTPA and EDTA neither terminated nor temporarily interrupted ICSI-induced Ca^2+^ oscillations (**Fig. 1C**), although protractedly slowed them down, possibly because of chelation and lowering of external Ca^2+^ (**Fig. 1C**). These results suggest that chelation of external Zn^2+^ does not affect the continuation of oscillations. We cannot determine that EDTA successfully chelated all external Zn^2+^, but the evidence that the addition of EDTA to the monitoring media containing cell impermeable FluoZin-3 caused a marked reduction in fluorescence, suggests that a noticeable fraction of the available Zn^2+^ was sequestered (**Supplementary Fig. 1A**). Similarly, injection of m*Plcζ* mRNA in eggs incubated in Ca^2+^ and Mg^2+^-free media supplemented with EDTA, to maximize the chances of chelation of external Zn^2+^, initiated low-frequency but persistent oscillations, and addition of Ca^2+^ and Mg^2+^ restored the physiological periodicity (**Supplementary Fig. 1B**). Lastly, another Zn^2+^-permeable chelator, TPA, blocked the ICSI-initiated Ca^2+^ oscillations but required higher concentrations than TPEN (**Fig. 1D**). Collectively, the data suggest that basal levels of labile internal Zn^2+^ are essential to sustain the fertilization-initiated Ca^2+^ oscillations in eggs.

**Figure 1.**
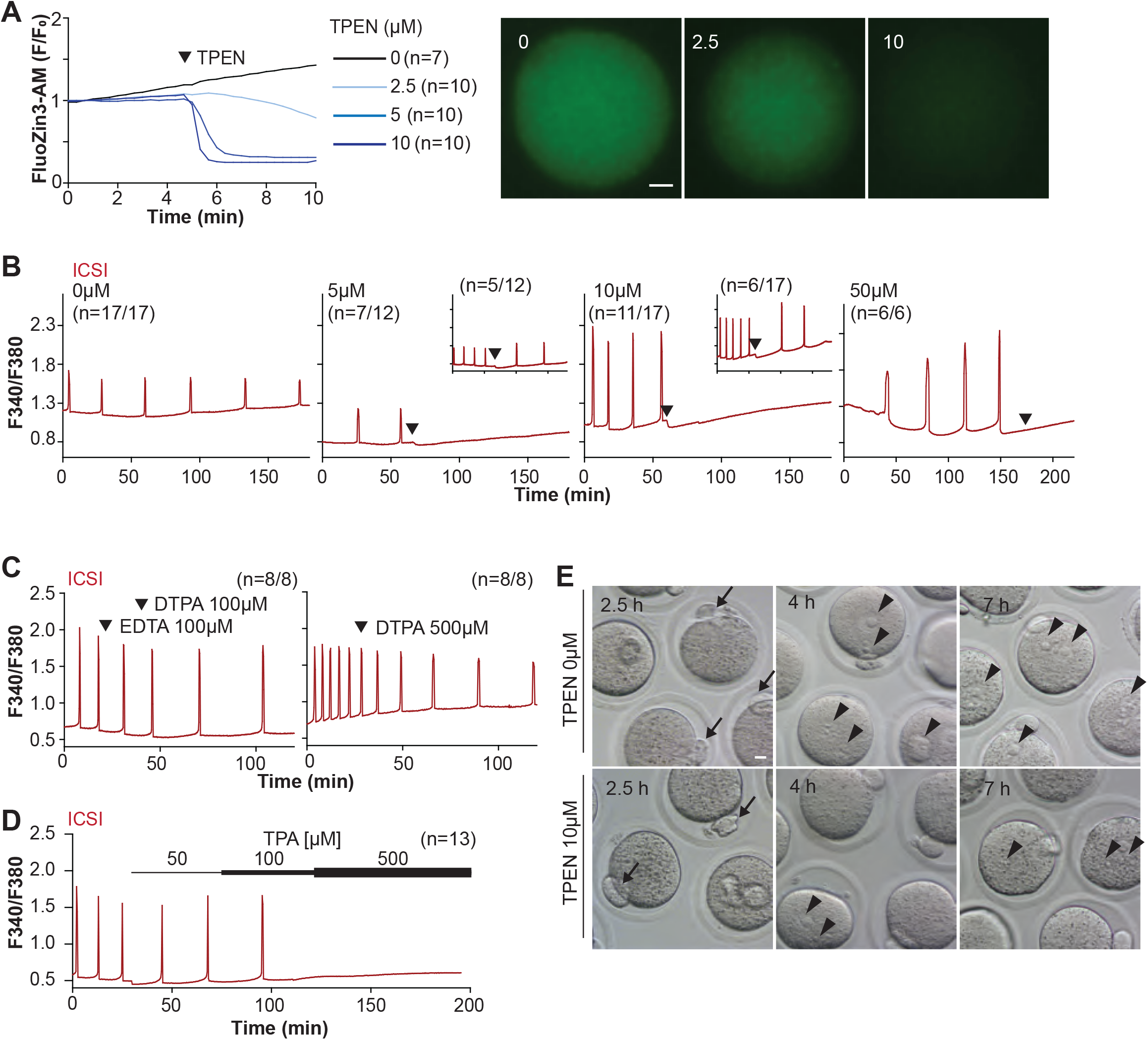
TPEN-induced Zn_2+_ deficiency inhibits fertilization-initiated Ca_2+_ oscillations in a dose-dependent manner. (**A**) (Left panel) Representative normalized Zn^2+^ recordings of MII eggs loaded with FluoZin-3AM following the addition of increasing concentrations of TPEN (0 µM, DMSO, black trace; 2.5 µM, sky blue; 5 µM, blue; 10 µM, navy). TPEN was directly added to the monitoring media. (Right panel) Representative fluorescent images of MII eggs loaded FluoZin-3AM supplemented with 0, 2.5, and 10 µM of TPEN. Scale bar: 10 µm. (**B-D**). (**B**) Representative Ca^2+^ oscillations following ICSI after the addition of 0, 5, 10, or 50 µM TPEN (arrowheads). Insets show representative traces for eggs that resumed Ca^2+^ oscillations after TPEN. (**C**) As above, but following the addition of 100 µM EDTA, 100 or 500 µM DTPA (time of addition denoted by arrowheads). (**D**) Ca^2+^ oscillations following ICSI after the addition of 50, 100, and 500 µM TPA (horizontal bars of increasing thickness). (**E**) Representative bright field images of ICSI fertilized eggs 2.5, 4, and 7 h after sperm injection. Arrows and arrowheads denote the second polar body and PN formation, respectively. Scale bar: 10 µm.

We next evaluated whether Zn^2+^ depletion prevented the completion of meiosis and pronuclear (PN) formation. To this end, ICSI-fertilized eggs were cultured in the presence of 10 μM TPEN for 8h, during which the events of egg activation were examined (**Fig. 1E and Table 1**). All fertilized eggs promptly extruded second polar bodies regardless of treatment (**Fig. 1E**). TPEN, however, impaired PN formation, and by 4- or 7-h post-ICSI, most treated eggs failed to show PNs, unlike controls (**Fig. 1E and Table 1**). Together, these results demonstrate that depletion of Zn^2+^ terminates Ca^2+^ oscillations and delays or prevents events of egg activation, including PN formation.

**Table 1.**
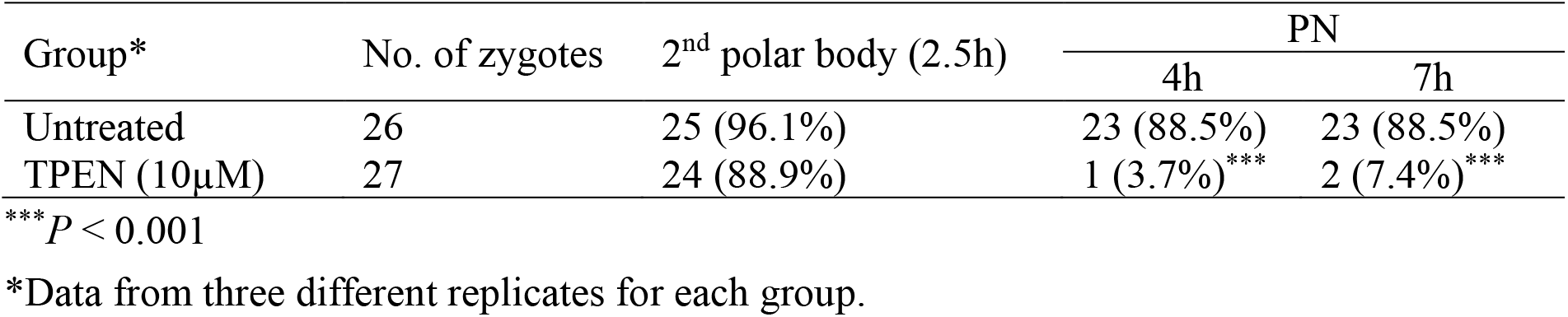
Addition of TPEN after ICSI does not prevent extrusion of the second polar body but precludes pronuclear (PN) formation.

### TPEN is a universal inhibitor of Ca_2+_ oscillations in eggs

Mammalian eggs initiate Ca^2+^ oscillations in response to numerous stimuli and conditions (Miyazaki and Ito, 2006; Wakai and Fissore, 2013). Fertilization and its release of PLCζ stimulate the phosphoinositide pathway, producing IP_3_ and Ca^2+^ oscillations (Miyazaki, 1988; Saunders et al., 2002). Neurotransmitters such as acetylcholine (Ach) and other G-protein coupled receptor agonists engage a similar mechanism (Dupont et al., 1996; Kang et al., 2003), although in these cases, IP_3_ production occurs at the plasma membrane and is short-lived (Kang et al., 2003; Swann and Parrington, 1999). Agonists such as SrCl_2_ and thimerosal generate oscillations by sensitizing IP_3_R1 without producing IP_3_. The mechanism(s) of SrCl_2_ is unclear, although its actions are reportedly directly on the IP_3_R1 (Hajnóczky and Thomas, 1997; Hamada et al., 2003; Nomikos et al., 2015, 2011; Sanders et al., 2018). Thimerosal oxidizes dozens of thiol groups in the receptor, which enhances the receptor’s sensitivity and ability to release Ca^2+^ (Bootman et al., 1992; Evellin et al., 2002; Joseph et al., 2018). We took advantage of the varied points at which the mentioned agonists engage the phosphoinositide pathway to examine TPEN’s effectiveness in inhibiting their effects. m*Plcζ* mRNA injection, like fertilization, induces persistent Ca^2+^ oscillations, although m*Plcζ*’s tends to be more robust. Consistent with this, the addition of 10 and 25 µM TPEN transiently interrupted or belatedly terminated oscillations, whereas 50 µM acutely stopped all responses (**Fig. 2A**). By contrast, SrCl_2_-initiated rises were the most sensitive to Zn^2+^-deficient conditions, with 2.5 µM TPEN nearly terminating all oscillations that 5 µM did (**Fig. 2B**). TPEN was equally effective in ending the Ach-induced Ca^2+^ responses (**Fig. 2C**), but curbing thimerosal responses required higher concentrations (**Fig. 2D**). Lastly, we ruled out that downregulation of IP_3_R1 was responsible for the slow-down or termination of the oscillations by TPEN. To accomplish this, we examined the IP_3_R1 mass in eggs (Jellerette et al., 2004) with and without TPEN supplementation and injection of m*Plc*ζ mRNA. By 4h post-injection, PLCζ induced the expected down-regulation of IP_3_R1 reactivity, whereas was insignificant in TPEN-treated and *Plc*ζ mRNA-injected eggs, as it was in uninjected control eggs (**Fig. 2F**). These findings together show that Zn^2+^ deficiency inhibits the IP_3_R1-mediated Ca^2+^ oscillations independently of IP_3_ production or loss of receptor, suggesting a role of Zn^2+^ on IP_3_R1 function (**Fig. 2E**).

**Figure 2.**
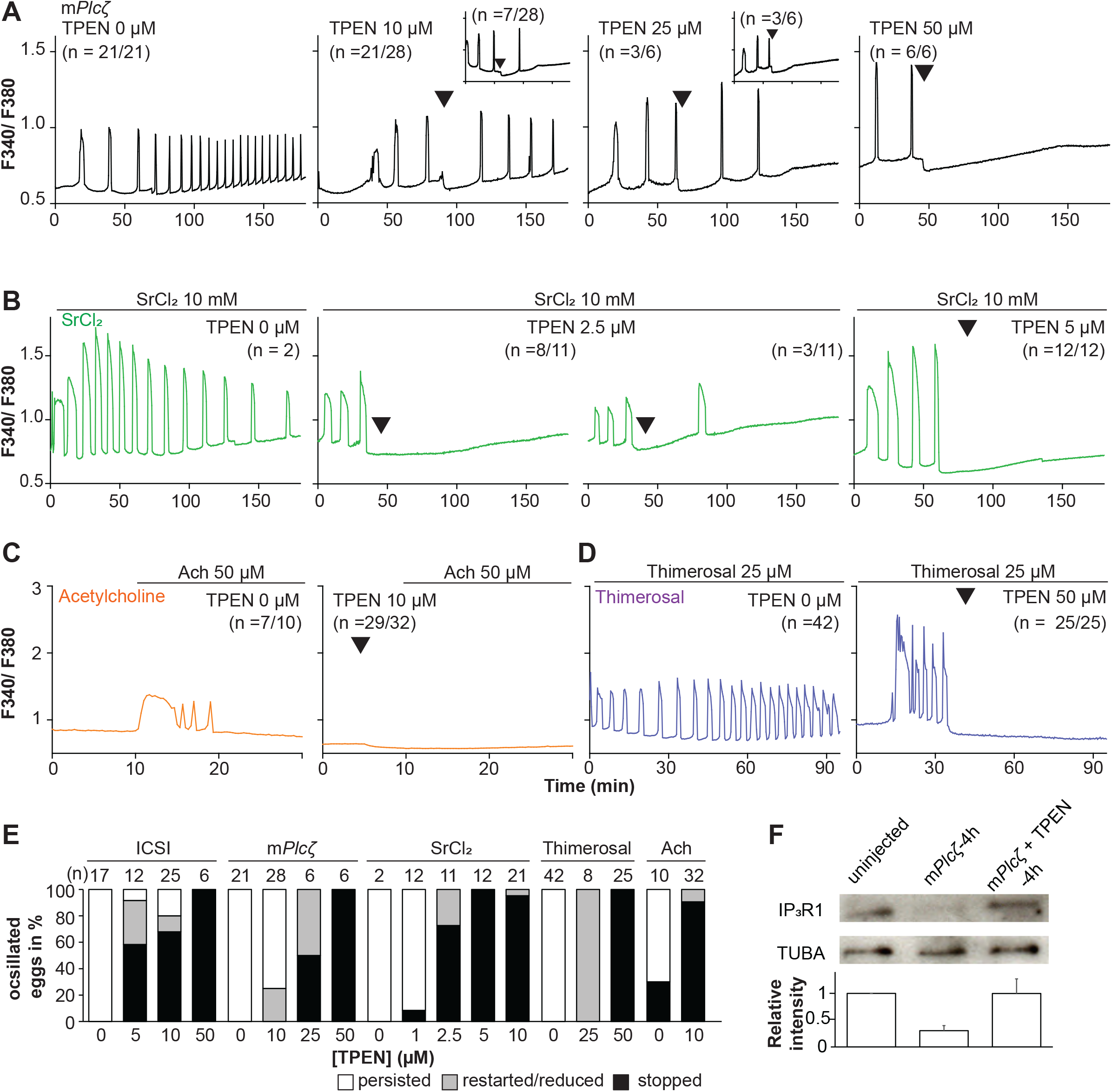
TPEN dose-dependently inhibits Ca_2+_ oscillations in eggs triggered by a broad-spectrum of agonists that stimulate the PI pathway or IP_3_R1. (**A-D**) Representative Ca^2+^ responses induced by (**A**) m*Plcζ* mRNA microinjection (0.01 µg/µl, black traces), (**B**) strontium chloride (10 mM, green), (**C**) acetylcholine chloride (50 µM, orange), and (**D**) thimerosal (25 µM, purple) in MII eggs. Increasing concentrations of TPEN were added to the monitoring media (arrowheads above traces denotes the time of adding). Insets in the upper row show representative traces of eggs that stop oscillating despite others continuing to oscillate. (**E**) Each bar graph summarizes the TPEN effect on Ca^2+^ oscillations at the selected concentrations for each of the agonists in A-D. (**F)** Western blot showing the intensities of IP_3_R1 and alpha-tubulin bands in MII eggs or in eggs injected with m*Plcζ* mRNA and incubated or not with TPEN above (*P* < 0.01). Thirty eggs per lane in all cases. This experiment was repeated twice, and the mean relative intensity of each blot is shown in the bar graph below.

### Zn^2+^ depletion reduces IP_3_R1-mediated Ca^2+^ release

To directly assess the inhibitory effects of TPEN on IP_3_R1 function, we used caged IP_3_ (cIP_3_) that, after short UV pulses, releases IP_3_ into the ooplasm (Wakai et al., 2012; Walker et al., 1987). To exclude the possible contribution of external Ca^2+^ to the responses, we performed the experiments in Ca^2+^-free media. In response to sequential cIP_3_ release 5 min apart, control eggs displayed corresponding Ca^2+^ rises that occasionally transitioned into short-lived oscillations (**Fig. 3A**). The addition of TPEN after the third cIP_3_ release prevented the subsequent Ca^2+^ response and prematurely terminated the in-progress Ca^2+^ rises (**Fig. 3B and inset)**. Pre-incubation of eggs with TPEN precluded cIP_3_-induced Ca^2+^ release, even after 5 sec UV exposure (**Fig. 3C**). The addition of excess ZnSO_4_ (100 µM) overcame TPEN’s inhibitory effects only if added before **(Fig. 3E)** and not after the addition of TPEN **(Fig. 3D)**. Similar concentrations of MgCl_2_ or CaCl_2_ failed to reverse TPEN effects **(Fig. 3F, G)**. Together, the results show that Zn^2+^ is required for IP_3_R1-mediated Ca^2+^ release downstream of IP_3_ production, appearing to interfere with receptor gating, as suggested by TPEN’s rapid termination of in-progress Ca^2+^ rises and ongoing oscillations.

**Figure 3.**
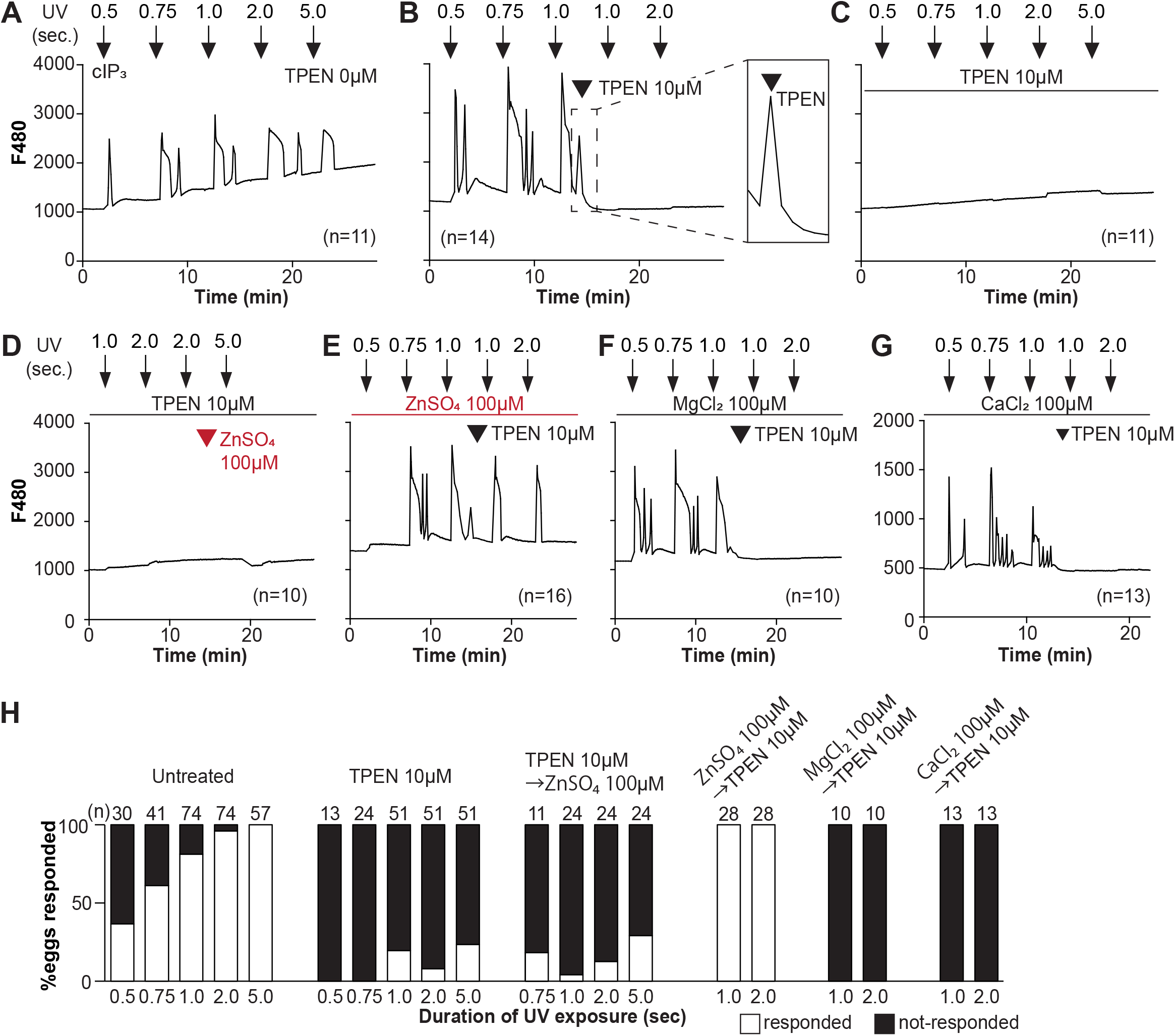
TPEN inhibition of cIP_3_-induced Ca^2+^ release is precluded by ZnSO_4_ supplementation before TPEN exposure. (**A-G**) Representative Ca^2+^ responses in MII eggs triggered by the release of caged IP_3_ (cIP_3_) induced by UV light pulses of increasing duration (arrows). (**A**) A representative control trace without TPEN, and (**B**) following the addition of 10 µM TPEN between the third and the fourth pulses. The broken line rectangle is magnified in the inset, farthest right side of the panel displaying the near immediate termination of an ongoing rise. (**C, D**) Recordings started in the presence of 10 µM TPEN but in (**D**) 100 µM ZnSO_4_ was added between the second and the third pulses. (**E**) Recording started in the presence of 100 µM ZnSO_4_ followed by the addition of 10 µM TPEN between the third and the fourth pulses. (**F, G**) Recording started in the presence of 100 µM MgSO_4_ (**F**) or 100 µM CaCl_2_ (**G**) and 10 µM TPEN added as above. Arrowheads above the different panels indicate the time of TPEN or divalent cation addition. (**H**) Bar graphs summarizing the number and percentages of eggs that responded to a given duration of UV pulses under each of the TPEN±divalent ions.

ERp44 is an ER luminal protein of the thioredoxin family that interacts with the IP_3_R1, reportedly inhibiting its ability to mediate Ca^2+^ release (Higo et al., 2005). The localization of ERp44 in the ER-Golgi intermediate compartment of somatic cells correlates with Zn^2+^’s availability and changes dramatically after TPEN treatment (Higo et al., 2005; Watanabe et al., 2019). To rule out the possibility that TPEN suppresses the function of IP_3_R1 by modifying the subcellular distribution of ERp44, we overexpressed ERp44 by injecting HA tagged-*Erp44* mRNA into MII eggs and monitored the effect on Ca^2+^ release. TPEN did not alter the localization of ERp44 (**Supplementary Fig. 2A**), and overexpression of ERp44 modified neither the Ca^2+^ oscillations induced by agonists (**Supplementary Fig. 2B**) nor the effectiveness of TPEN to block them (data not shown). Thus, TPEN and Zn^2+^ deficiency most likely inhibits Ca^2+^ release by directly interfering with IP_3_R1 function rather than modifying this particular regulator.

### Zn_2+_ depletion diminishes the ER Ca_2+_ leak and increases Ca_2+_ store content

Our above cIP_3_ results that TPEN inhibited IP_3_R1-mediated Ca^2+^ release and interrupted in-progress Ca^2+^ rises despite the presence of high levels of environmental IP_3_ suggest its actions are probably independent of IP_3_ binding, agreeing with an earlier report showing that TPEN did not modify IP_3_’s affinity for the IP_3_R (Richardson and Taylor, 1993). Additionally, the presence of a Zn^2+^-binding motif near the C-term cytoplasmic domain of the IP_3_R1’s channel, which is known to influence agonist-induced IP_3_R1 gating (Fan et al., 2015), led us to posit and examine that Zn^2+^ deficiency may be disturbing Ca^2+^ release to the cytosol and out of the ER. To probe this possibility, we queried if pre-treatment with TPEN inhibited Ca^2+^ release through IP_3_R1. We first used Thapsigargin (Tg), a Sarcoplasmic/ER Ca^2+^ ATPase pump inhibitor (Thastrup et al., 1990) that unmasks a constitutive Ca^2+^ leak out of the ER (Lemos et al., 2021); in eggs, we have demonstrated it is mediated at least in part by IP_3_R1 (Wakai et al., 2019). Treatment with TPEN for 15 min slowed the Tg-induced Ca^2+^ leak into the cytosol, resulting in delayed and lowered amplitude Ca^2+^ responses (**Fig. 4A**; *P*<0.05). To test whether the reduced response to Tg means that TPEN prevented the complete response of Tg, leaving a temporarily increased Ca^2+^ content in the ER, we added the Ca^2+^ ionophore ionomycin (Io), which empties all stores independently of IP_3_Rs. Io-induced Ca^2+^ responses were 3.3-fold greater in TPEN-treated cells, supporting the view that TPEN interferes with the ER Ca^2+^ leak **(Fig. 4A**; *P* < 0.05). We further evaluated this concept using *in vitro* aged eggs that often display reduced Ca^2+^ store content than freshly collected counterparts (Abbott et al., 1998). After culturing eggs in the presence or absence of TPEN for 2h, we added Io during Ca^2+^ monitoring, which in TPEN-treated eggs induced bigger Ca^2+^ rises than in control eggs **(Fig. 4B**; *P* <0.05). We confirmed that this effect was independent of IP_3_R1 degradation because TPEN did not change IP_3_R1 reactivity in unfertilized eggs (**Fig. 4C**; *P* <0.05).

**Figure 4.**
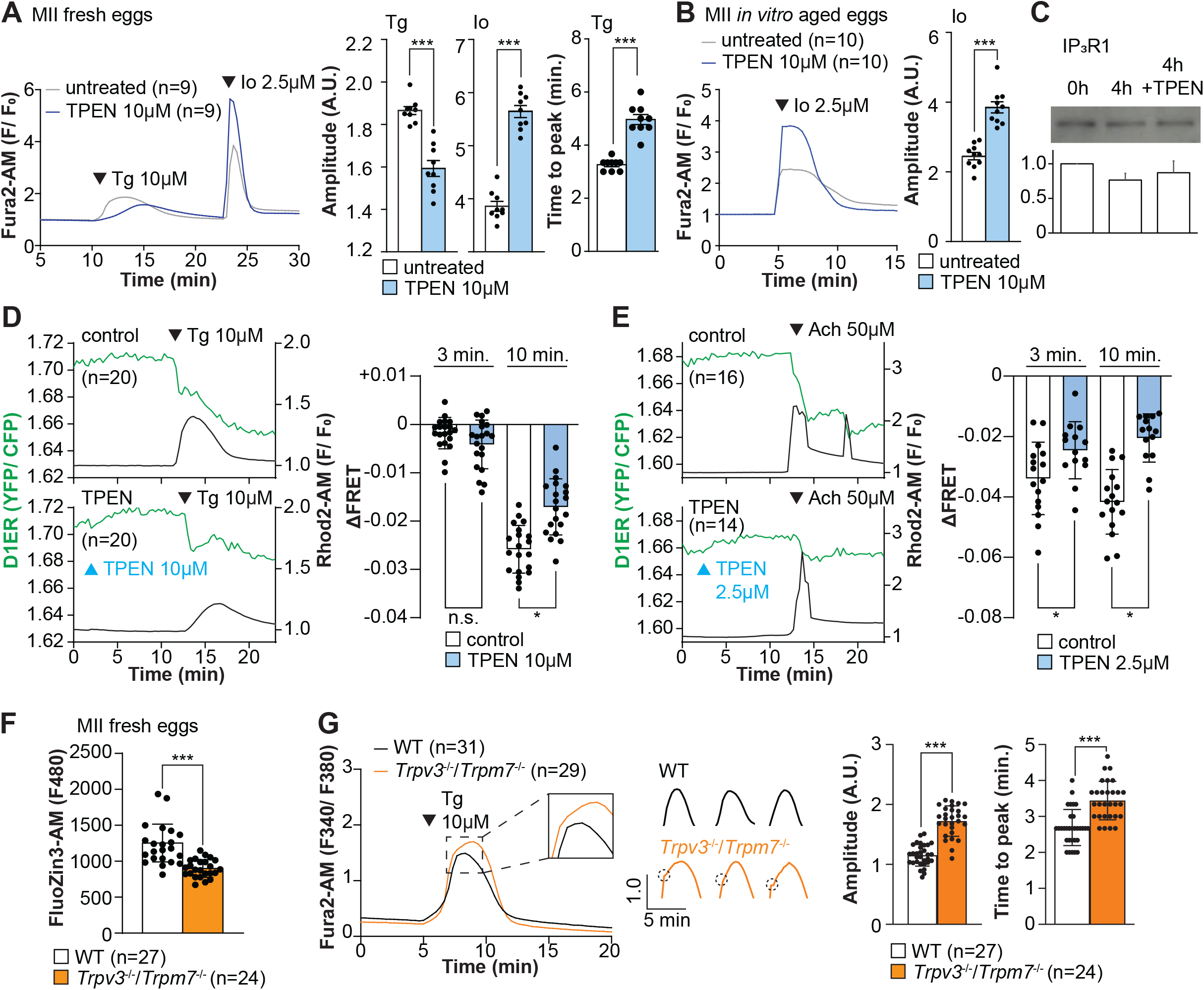
Zn_2+_ depletion alters Ca_2+_ homeostasis and increases Ca_2+_ store content independent of IP_3_R1 mass. (**A, B**) Representative Ca^2+^ traces of MII eggs after the addition of Tg and Io in the presence or absence of TPEN. Blue trace recordings represent TPEN-treated eggs whereas gray traces represent control, untreated eggs. (**A**) Io was added to fresh MII eggs once Ca^2+^ returned to baseline after treatment with Tg. Comparisons of mean peak amplitudes after Tg and Io are shown in the bar graphs in the right panel (*P* < 0.001). (**B**) MII eggs were aged by 2h. incubation supplemented or not with TPEN followed by Io addition and Ca^2+^ monitoring (*P* < 0.001). (**C)** Western blot showing the intensities of IP_3_R1 bands in MII eggs freshly collected, aged by 4h. incubation without TPEN, and with TPEN. Thirty eggs per lane in all cases. This experiment was repeated three times. (**D, E**) (Left panels) Representative traces of Ca^2+^ values in eggs loaded with the Ca^2+^-sensitive dye Rhod-2 AM and the ER Ca^2+^reporter, D1ER (1 µg/µl mRNA). TPEN was added into the media followed 10 min later by (**D**) 10 µM Tg and (**E**) 50 µM Ach. (Right panel) Bars represent the difference of FRET value between at the time of Tg/ Ach addition and at 3 and 5 min later of the addition (*P* < 0.05). Experiments were repeated two different times for each treatment. Black and green traces represent cytosolic Ca^2+^ and Ca^2+^-ER, respectively. Blue and black arrowheads indicate the time of addition of TPEN and Tg/ Ach, respectively. (**F**) Basal Zn^2+^ level comparison in WT (open bar) and *Trpv3^-/-^*/*Trpm7^-/-^*(dKO, orange bar) MII eggs. Each plot represents the Fluozin3 measurement at 5 min after starting monitoring. (**G**) (Left panel) Representative Ca^2+^ traces of WT (black trace) and dKO (orange trace) MII eggs after adding Tg. Insets represent the magnified traces at the peak of Ca^2+^ spike from different sets of eggs. (Middle panel) Individual traces of WT and dKO eggs after Tg addition. Dashed circles represent the flection point in dKO traces. (Right panel) Comparisons of mean peak amplitudes after Tg and the time between Tg addition and the Ca^2+^ peak are shown in the bar graphs in the right panel (*P* < 0.001).

Next, we used the genetically encoded FRET sensor D1ER (Palmer et al., 2004) to assess the TPEN’s effect on the ER’s relative Ca^2+^ levels changes following the additions of Tg or Ach. TPEN was added 10 min before 10 µM Tg or 50 µM Ach, and we simultaneously monitored changes in cytosolic and intra-ER Ca^2+^ (**Fig. 4D, E**). For the first three min, the Tg-induced decrease in Ca^2+^-ER was similar between groups. However, while the drop in Ca^2+^ content continued in control eggs, in TPEN-treated eggs, it came to an abrupt halt, generating profound differences between the two groups (**Fig. 4D**; *P* <0.05). TPEN had even more pronounced effects following the addition of Ach, leading to a reduced- and prematurely terminated-Ca^2+^ release from the ER in treated eggs (**Fig. 4E**; *P* <0.05).

Lastly, we sought to use a cellular model where low labile Zn^2+^ occurred without pharmacology. To this end, we examined a genetic model where the two non-selective plasma membrane channels that could influx Zn^2+^ in maturing oocytes have been deleted (Bernhardt et al., 2017; Carvacho et al., 2016, 2013), namely, the transient receptor potential melastatin-7 (TRPM7) and TRP vanilloid 3 (TRPV3), both members of the TRP superfamily of channels (Wu et al., 2010). We found that eggs from double knockout females (dKOs) had lower levels of labile Zn^2+^ (**Fig. 4F**), and the addition of Tg revealed an expanded Ca^2+^ store content in these eggs vs. control WT eggs (**Fig. 4G**). Remarkably, in dKO eggs, the Ca^2+^ rise induced by Tg showed a shoulder or inflection point before the peak delaying the time to peak (**Fig. 4G**, **inset**; *P* <0.001). These results in dKO eggs show a changed dynamic of the Tg-induced Ca^2+^ release, suggesting that lower levels of labile Zn^2+^ modify ER Ca^2+^ release independently of chelators.

### Ca_2+_ oscillations in eggs occur within a window of Zn_2+_ concentrations

We next examined if resupplying Zn^2+^ could restart the Ca^2+^ oscillations terminated by Zn^2+^ depletion. Zn pyrithione (ZnPT) rapidly increases cellular Zn^2+^ upon extracellular addition (Barnett et al., 1977; Robinson, 1964). Dose titration studies and imaging fluorimetry revealed that 0.01 µM ZnPT caused subtle and protracted increases in Zn^2+^ levels, whereas 0.1 µM ZnPT caused rapid increases in eggs’ Zn^2+^ baseline (**Fig. 5A**). We induced detectable Ca^2+^ oscillations by injection of m*Plc*ζ mRNA followed by 50 µM TPEN (**Fig. 5B**), which terminated them. After 30 min, we added 0.1 µM ZnPT, and within 15 min the oscillations restarted in most TPEN-treated eggs (**Fig. 5C**). We repeated this approach using Thimerosal (**Fig. 5D, E**). Adding 0.1 µM ZnPT did not restore the Ca^2+^ oscillations retrained by TPEN, but 0.5 µM ZnPT did so (**Fig. 5E**). These results demonstrate that Zn^2+^ plays a pivotal, enabling role in the generation of Ca^2+^ oscillations in mouse eggs.

**Figure 5.**
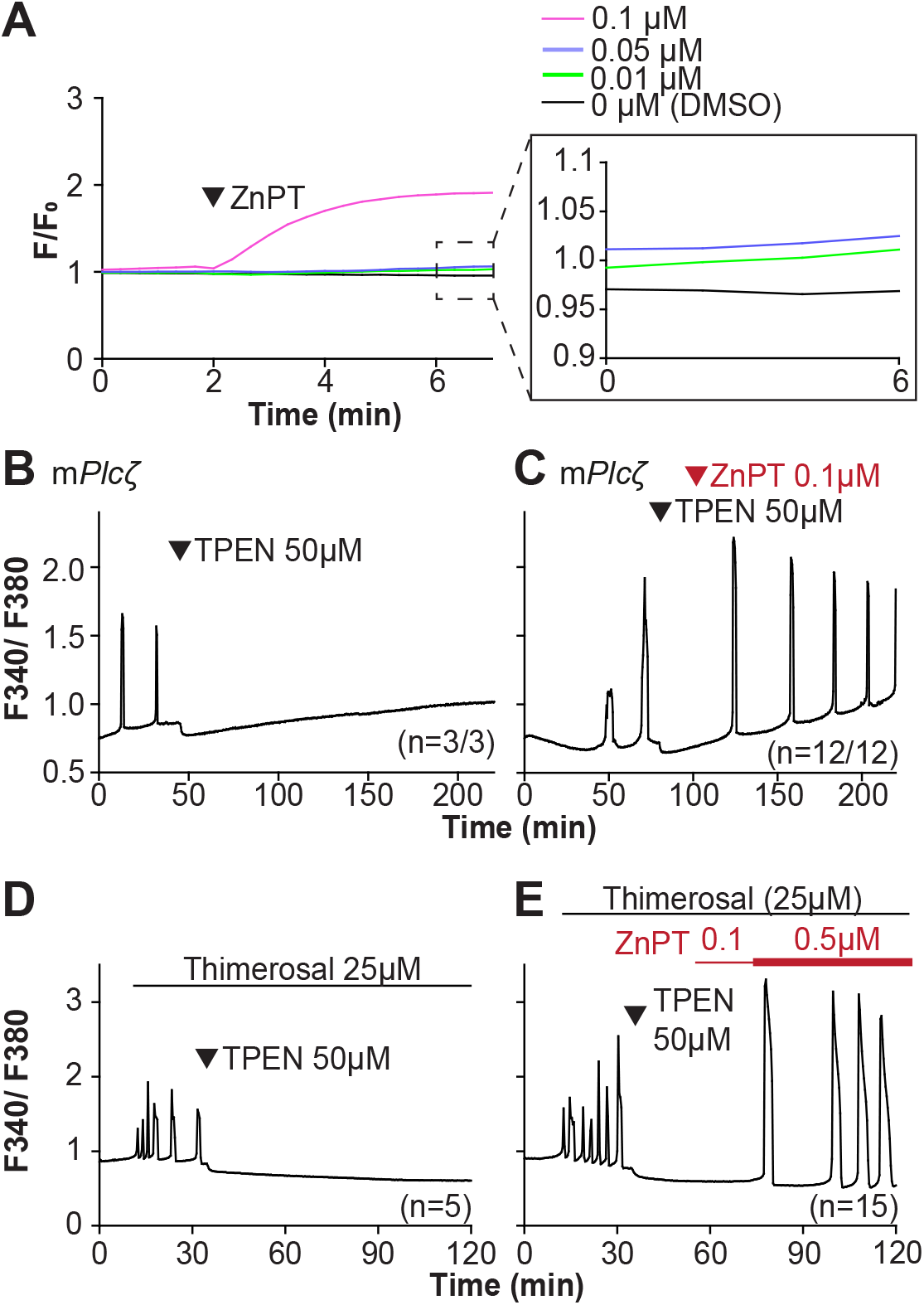
Restoring Zn_2+_ levels with ZnPT rescues oscillations interrupted by TPEN-induced Zn_2+_ deficiency. (**A**) Representative traces of Zn^2+^ in MII eggs following the addition of 0.01 to 0.1 µM concentrations of ZnPT. The broken rectangular area is amplified in the next panel to appreciate the subtle increase in basal Zn^2+^ caused by the addition of ZnPT. (**B, C**) m*Plcζ* mRNA (0.01 µg/µl)-induced oscillations followed by the addition of TPEN (black arrowhead) (**B**), or after the addition of TPEN followed by ZnPT (red arrowhead) (**C**). (**D, E**) Thimerosal (25 µM) induced oscillations using the same sequence of TPEN (**D**) and ZnPT (**E**), but higher concentrations of ZnPT were required to rescue Thimerosal-initiated oscillations (**E**). These experiments were repeated at least two different times.

### Excessive intracellular Zn_2+_ inhibits Ca_2+_ oscillations

Zn^2+^ is necessary for diverse cellular functions, consistent with numerous amino acids and proteins capable of binding Zn^2+^ within specific and physiological ranges (Pace and Weerapana, 2014). Excessive Zn^2+^, however, can cause detrimental effects on cells and organisms (Broun et al., 1990; Hara et al., 2022; Sikora and Ouagazzal, 2021). Consistent with the deleterious effects of Zn^2+^, a previous study showed that high concentrations of ZnPT, ∼50 µM, prevented SrCl_2_-induced egg activation and initiation of development (Bernhardt et al., 2012; Kim et al., 2011). We examined how ZnPT and excessive Zn^2+^ levels influence Ca^2+^ oscillations. Our conditions revealed that pre-incubation or continuous exposure to 0.1 µM or 1.0 µM ZnPT delayed or prevented egg activation induced by m*Plc*ζ mRNA injection (**Supplementary Fig. 3**). We used these ZnPT concentrations to add it into ongoing oscillations induced by ICSI and monitored the succeeding Ca^2+^ responses. The addition of 0.05 to 10 µM ZnPT caused an immediate elevation of the basal levels of Fura-2 and termination of the Ca^2+^ oscillations (**Fig. 6A-D)**. m*Plc*ζ mRNA-initiated Ca^2+^ responses were also interrupted by adding 0.1 µM ZnPT, whereas untreated eggs continued oscillating (**Fig. 6E, F**). ZnPT also inhibited IP_3_R1-mediated Ca^2+^ release triggered by cIP_3_, suggesting that excessive Zn^2+^ directly inhibits IP_3_R1 function (**Fig. 6G)**.

**Figure 6.**
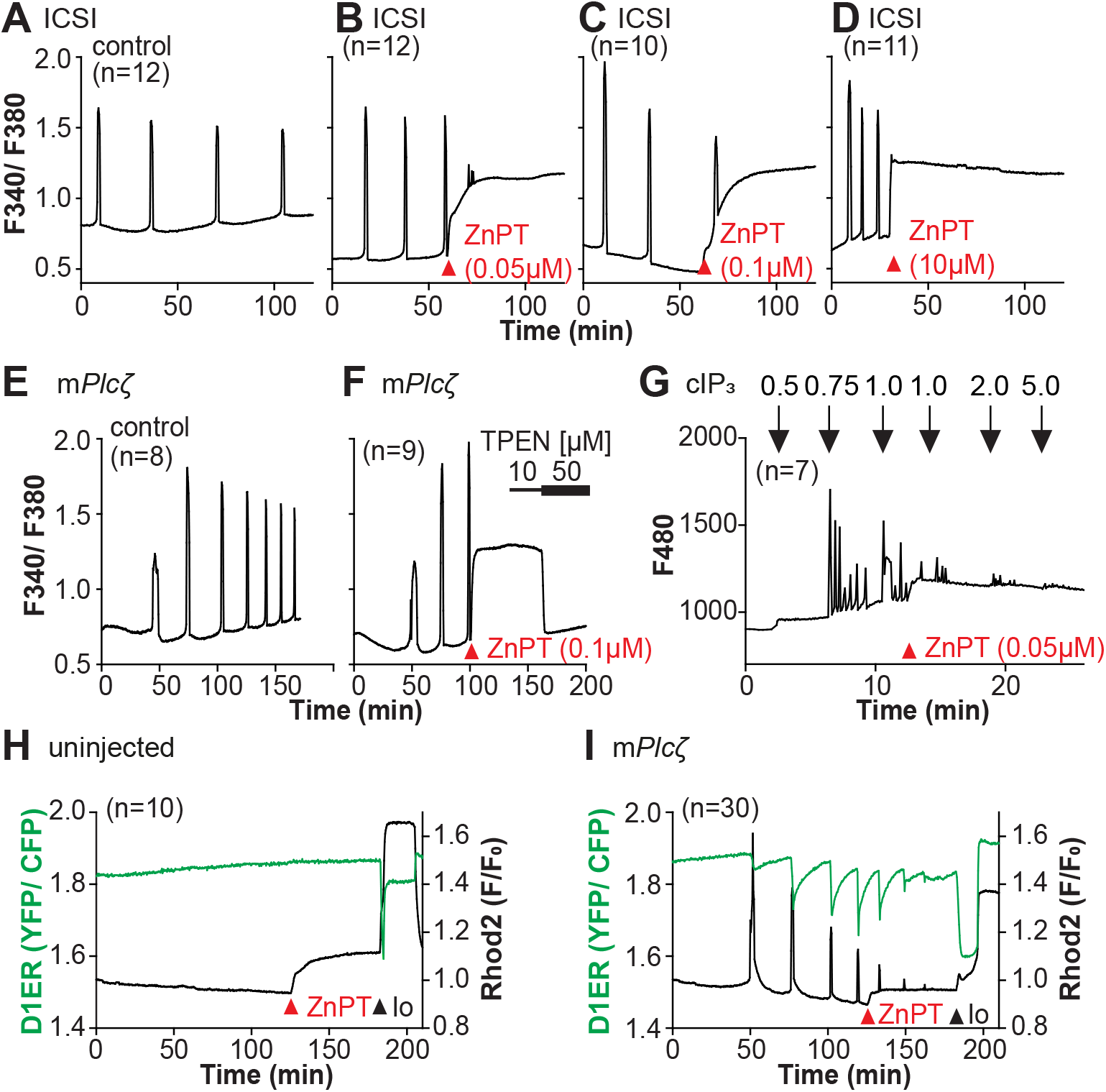
Excess Zn_2+_ hinders Ca_2+_ oscillations. (**A-D**) ICSI-initiated Ca^2+^ response without (**A**) or following the addition of ZnPT (**B, C**) (the time of ZnPT addition and concentration are denoted above the tracing). (**E**, **F**) Representative Ca^2+^ responses induced by injection of 0.01 µg/µl m*Plcζ* mRNA in untreated eggs (**E**) or in eggs treated with 0.1 µM ZnPT followed by 10 µM TPEN first and then 50 µM (**F**). (**G**) cIP_3_-induced Ca^2+^ release as expected when the UV pulses in the absence but not in the presence of 0.05 µM ZnPT (the time of addition is denoted by a bar above the tracing). (**H, I**) Representative traces of Ca^2+^ values in eggs loaded with the Ca^2+^-sensitive dye Rhod-2 AM and the ER Ca^2+^reporter, D1ER (1 µg/µl mRNA). Uninjected and 0.01 µg/µl m*Plcζ* mRNA-injected eggs were monitored. After initiation and establishment of the oscillations, 0.1 µM ZnPT was added into the media followed 30 min later by 2.5 µM Io. Experiments were repeated two different times. Red and black arrowheads indicate the time of addition of ZnPT and Io, respectively.

A noticeable feature of ZnPT is the increased basal ratios of Fura-2 AM. These changes could reflect enhanced IP_3_R1 function and increased basal Ca^2+^ concentrations caused by Zn^2+^ stimulation of IP_3_R1. This seems unlikely, however, because extended elevated cytosolic Ca^2+^ would probably induce cellular responses, such as the release of the second polar body, egg fragmentation, or cell death, neither of which happened. It might reflect, instead, Fura-2’s ability to report changes in Zn^2+^ levels, which seemed the case because the addition of TPEN lowered fluorescence without restarting the Ca^2+^ oscillations (**Fig. 6F**). To ensure the impact of ZnPT abolishing Ca^2+^ oscillations was not an imaging artifact obscuring ongoing rises, we simultaneously monitored cytoplasmic and ER Ca^2+^ levels with Rhod-2 and D1ER, respectively. This approach allowed synchronously observing Ca^2+^ changes in both compartments that should unfold in opposite directions. In control, uninjected eggs, the fluorescent values for both reporters remained unchanged during the monitoring period, whereas in m*Plc*ζ mRNA-injected eggs, the reporters’ signals displayed simultaneous but opposite changes, as expected (**Fig. 6H, I**). The addition of ZnPT in uninjected eggs rapidly increased Rhod-2 signals but not D1ER’s, which was also the case in oscillating eggs, as the addition of ZnPT did not immediately alter the dynamics of the ER’s Ca^2+^ release, suggesting D1ER faithfully reports in Ca^2+^ changes but cannot detect changes in Zn^2+^ levels, at least to this extent; ZnPT progressively caused fewer and lower amplitude changes in D1ER fluorescence, consistent with the diminishing and eventual termination of the Ca^2+^ oscillations. Noteworthy, in these eggs, the basal D1ER fluorescent ratio remained unchanged after ZnPT, demonstrating its unresponsiveness to Zn^2+^ changes of this magnitude. The ZnPT-induced increases in Rhod-2 fluorescence without concomitant changes in D1ER values suggest that the changes in the dyes’ fluorescence do not represent an increase in basal Ca^2+^ and, more likely, signal an increase in intracellular Zn^2+^. We confirmed that both reporters were still in working order, as the addition of Io triggered Ca^2+^ changes detected by both reporters (**Fig. 6H, I**).

## Discussion

The present study demonstrates that appropriate levels of labile Zn^2+^ are essential for initiating and maintaining IP_3_R1-mediated Ca^2+^ oscillations in mouse eggs regardless of the initiating stimuli. Both deficient and excessive Zn^2+^ compromise IP_3_R1 sensitivity, diminishing and mostly terminating Ca^2+^ oscillations. The results demonstrate that IP_3_R1 and Zn^2+^ act in concert to modulate Ca^2+^ signals, revealing previously unexplored crosstalk between these ions at fertilization (**Fig. 7**).

**Figure 7.**
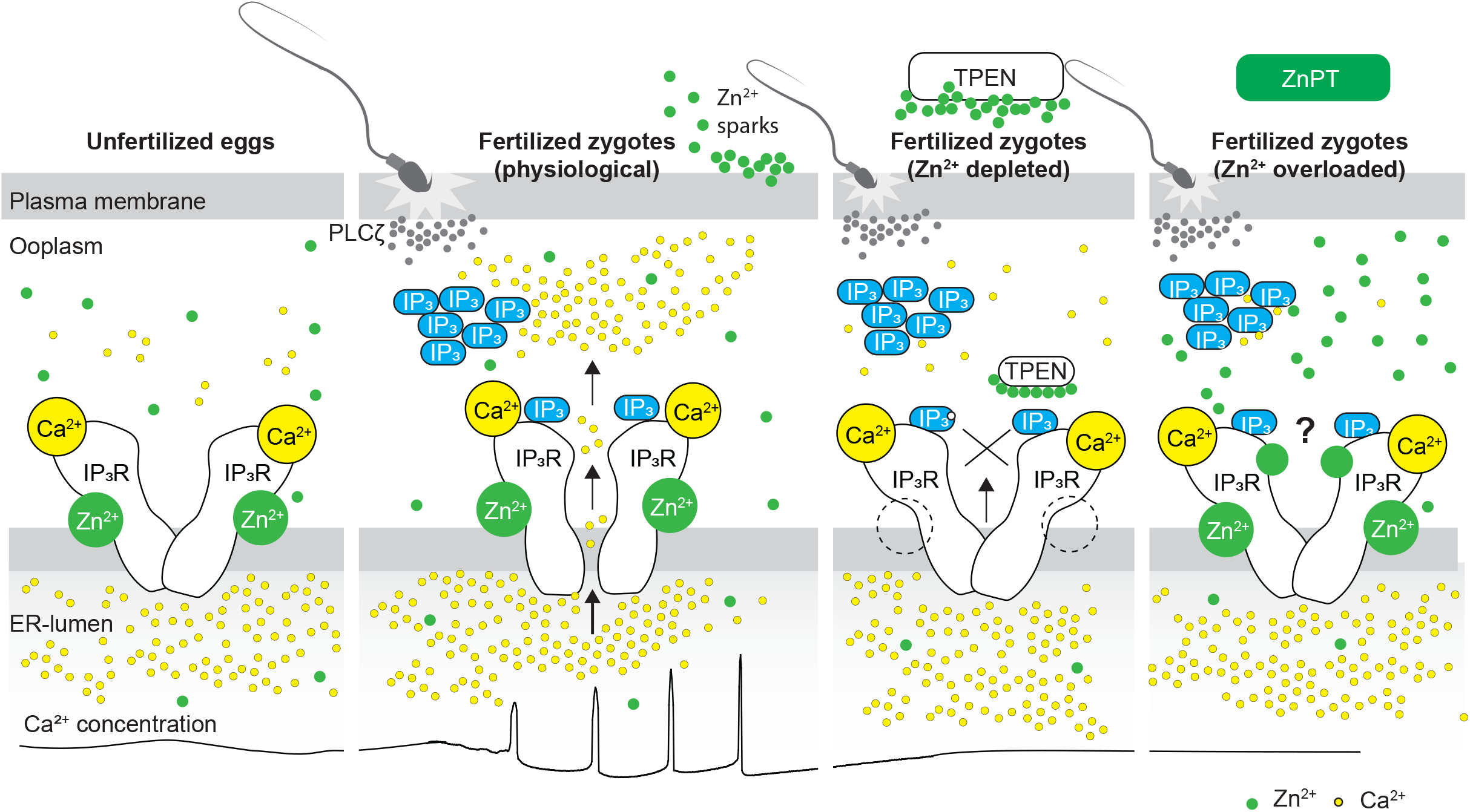
Schematic of proposed regulation of IP_3_R1 function by Zn^2+^ in eggs and fertilized zygotes. In MII eggs, left panel, IP_3_R1s are in a Ca^2+^-release permissive state with optimal levels of cytoplasmic Ca^2+^ and Zn^2+^ and maximum ER content, but Ca^2+^ is maintained at resting levels by the combined actions of pumps, ER Ca^2+^ leak, and reduced influx. Once fertilization takes place, left center panel, robust IP_3_ production induced by the sperm-borne PLCζ leads to Ca^2+^ release through ligand-induced gating of IP_3_R1. Continuous IP_3_ production and refilling of the stores via Ca^2+^ influx ensure the persistence of the oscillations. Zn^2+^ release occurs in association with first few Ca^2+^ rises and cortical granule exocytosis, Zn^2+^ sparks, lowering Zn^2+^ levels but not sufficiently to inhibit IP_3_R1 function. Zn^2+^ deficiency caused by TPEN or other permeable Zn^2+^ chelators, right center panel, dose-dependently impairs IP_3_R1 function and limits Ca^2+^ release. We propose this is accomplished by stripping the Zn^2+^ bound to the residues of the zinc-finger motif in the LNK domain of IP_3_R1 that prevents the allosteric modulation of the gating process induced by IP_3_ or other agonists. We propose that excess Zn^2+^, right panel, also inhibits IP_3_R1-mediate Ca^2+^ release, possibly by non-specific binding of thiol groups present in cysteine residues throughout the receptor (denoted by a ?). We submit that optimal Ca^2+^ oscillations in mouse eggs unfold in the presence of a permissive range of Zn^2+^ concentration.

Zn^2+^ is an essential micronutrient for living organisms (Kaur et al., 2014) and is required for various cellular functions, such as proliferation, transcription, and metabolism (Lo et al., 2020; Maret and Li, 2009; Yamasaki et al., 2007). Studies using Zn^2+^ chelators have uncovered what appears to be a cell-specific, narrow window of Zn^2+^ concentrations needed for cellular proliferation and survival (Carraway and Dobner, 2012; Lo et al., 2020). Further, TPEN appeared especially harmful, and in a few cell lines, even low doses provoked oxidative stress, DNA fragmentation, and apoptosis (Mendivil-Perez et al., 2012). We show here that none of the Zn^2+^ chelators, permeable or impermeable, affected cell viability within our experimental observations, confirming findings from previous studies that employed high concentrations of TPEN to interrupt the Ca^2+^ oscillations (Lawrence et al., 1998) or inducing egg activation of mouse eggs (Suzuki et al., 2010b). Our data demonstrating that ∼2.5 µM is the threshold concentration of TPEN in eggs that first causes noticeable changes in basal Zn^2+^, as revealed by FluoZin, is consistent with the ∼2 to 5-µM Zn^2+^ concentrations in most culture media without serum supplementation (Lo et al., 2020), and with the ∼100 pM basal Zn^2+^ in cells (Qin et al., 2011). Lastly, the effects on Ca^2+^ release observed here with TPEN and other chelators were due to the chelation of Zn^2+^, as pretreatment with ZnSO_4_ but not with equal or greater concentrations of MgCl_2_ or CaCl_2_ rescued the inhibition of the responses, which is consistent with results by others (Kim et al., 2010; Lawrence et al., 1998).

To identify how Zn^2+^ deficiency inhibits Ca^2+^ release in eggs, we induced Ca^2+^ oscillations using various stimuli and tested the effectiveness of membrane-permeable and impermeable chelators to abrogate them. Chelation of extracellular Zn^2+^ failed to terminate the Ca^2+^ responses, whereas membrane-permeable chelators did, pointing to intracellular labile Zn^2+^ levels as essential for Ca^2+^ release. All agonists used here were susceptible to inhibition by TPEN, whether their activities depended on IP_3_ production or allosterically induced receptor function, although the effective TPEN concentrations varied across stimuli. Some agents, such as m*Plc*ζ mRNA or thimerosal, required higher concentrations than SrCl_2_, Ach, or cIP_3_. The reason underlying the different agonists’ sensitivities to TPEN will require additional research, but the persistence of IP_3_ production or change in IP_3_R1 structure needed to induce channel gating might explain it. However, the universal abrogation of Ca^2+^ oscillations by TPEN supports the view drawn from cryo-EM-derived IP_3_R1 models that signaling molecules can allosterically induce channel gating from different starting positions in the receptor by mechanically coupling the binding effect to the ion-conducting pore in the C-terminal end of IP_3_R (Fan et al., 2015). The cytosolic C-terminal domain of each IP_3_R1 subunit is alongside the IP_3_-binding domain of another subunit and, therefore, well positioned to sense IP_3_ binding and induce channel gating (Fan et al., 2015). Within each subunit, the LNK domain, which contains a Zn^2+^-finger motif (Fan et al., 2015), connects the opposite domains of the molecule. Although there are no reports regarding the regulation of IP_3_R1 sensitivity by Zn^2+^, such evidence exists for RyRs (Woodier et al., 2015), which also display a conserved Zn^2+^-finger motif (des Georges et al., 2016). Lastly, mutations of the two Cys or two His residues of this motif, without exception, resulted in inhibition or inactivation of the IP_3_R1 channel (Bhanumathy et al., 2012; Uchida et al., 2003). These results are consistent with the view that the C-terminal end of IP_3_Rs plays a dominant role in channel gating (Bhanumathy et al., 2012; Uchida et al., 2003). We propose that TPEN inhibits Ca^2+^ oscillations in mouse eggs because chelating Zn^2+^ interferes with the function of the LNK domain and its Zn^2+^-finger motif proposed role on the mechanical coupling induced by agonist binding to the receptor that propagates to the pore-forming region and required to gate the channel’s ion-pore (Fan et al., 2022, 2015).

In support of this possibility, TPEN-induced Zn^2+^ deficient conditions altered the Ca^2+^-releasing kinetics in resting eggs or after fertilization. Tg increases intracellular Ca^2+^ by inhibiting the SERCA pump (Thastrup et al., 1990) and preventing the reuptake into the ER of the ebbing Ca^2+^ during the basal leak. Our previous studies showed that the downregulation of IP_3_R1 diminishes the leak, suggesting it occurs through IP_3_R1 (Wakai and Fissore, 2019). Consistent with this view, TPEN pre-treatment delayed the Ca^2+^ response induced by Tg, implying that Zn^2+^ deficiency hinders Ca^2+^ release through IP_3_R1. An expected consequence would be increased Ca^2+^ content in the ER after Tg. Io that mobilizes Ca^2+^ independently of IP_3_Rs (Toeplitz et al., 1979) induced enhanced responses in TPEN-treated eggs vs. controls, confirming the accumulation of Ca^2+^-ER in Zn^2+^ deficient conditions. We demonstrated that this accumulation is due to hindered emptying of the Ca^2+^ ER evoked by agonists in Zn^2+^-deficient environments, resulting in reduced cytosolic Ca^2+^ increases, as IP_3_R1 is the pivotal intermediary channel between these compartments. Noteworthy, the initial phase of the Tg-induced Ca^2+^ release out of the ER did not appear modified by TPEN, as if it was mediated by a Zn^2+^-insensitive Ca^2+^ channel(s)/transporter, contrasting with the abrogation of Ach-induced ER emptying from the outset. Remarkably, independently of Zn^2+^ chelators, emptying of Ca^2+^ ER was modified in a genetic model of Zn^2+^-deficient oocytes lacking two TRP channels, confirming the impact of Zn^2+^ on Ca^2+^ release. It is worth noting that TPEN did not reduce but maintained or increased the mass of IP_3_R1, which might result in the inhibition of Zn^2+^-dependent ubiquitin ligase Ubc7 by the Zn-deficient conditions (Webster et al., 2003). We cannot rule out that these conditions may undermine other conformational changes required to trigger IP_3_R1 degradation, thereby favoring the accumulation of IP_3_R1.

Despite accruing Zn^2+^ during oocyte maturation, fertilization witnesses a necessary Zn^2+^ release into the external milieu, known as “Zn^2+^ sparks” (Converse and Thomas, 2020; Kim et al., 2011; Mendoza et al., 2022; Que et al., 2019, 2015; Seeler et al., 2021). This release of Zn^2+^ is a conserved event in fertilization across species and is associated with several biological functions, including those related to fending off polyspermy (Kim et al., 2011; Que et al., 2019; Wozniak et al., 2020). The concomitant decrease in Zn^2+^ facilitates the resumption of the cell cycle and exit from the MII stage (Kim et al., 2011). Congruent with this observation, artificial manipulation that maintains high Zn^2+^ levels prevents egg activation (Kim et al., 2011), whereas lowering Zn^2+^ with chelators leads to egg activation without Ca^2+^ mobilization (Suzuki et al., 2010b). As posed by others, these results suggest that meiosis completion and the early stages of fertilization unfold within a narrow window of permissible Zn^2+^ (Kim et al., 2011, 2010). Here, we extend this concept and show that IP_3_R1 function and the Ca^2+^ oscillations in mouse eggs require this optimal level of labile Zn^2+^ because the Ca^2+^ responses interrupted by TPEN-induced Zn^2+^-insufficiency are rescued by restoring Zn^2+^ levels with ZnPT. Furthermore, unopposed increases in Zn^2+^ by exposure to ZnPT abrogated fertilization-initiated Ca^2+^ oscillations and prevented the expected egg activation events. It is unclear how excess Zn^2+^ disturbs the function of IP_3_R1. Nevertheless, IP_3_R1s have multiple cysteines whose oxidation enhances the receptor sensitivity to IP_3_ (Joseph et al., 2018), and it is possible that excessive Zn^2+^ aberrantly modifies them, disturbing IP_3_R1 structure and function or, alternatively, preventing their oxidation and sensitization of the receptor. Lastly, we cannot rule out that high Zn^2+^ levels directly inhibit the receptor’s channel. These results reveal a close association between the Zn^2+^ levels controlling meiotic transitions and the Ca^2+^ release necessary for egg activation, placing the IP_3_R1 at the center of the crosstalk of these two divalent cations.

Abrupt Zn^2+^ changes have emerged as critical signals for meiotic and mitotic transitions in oocytes, eggs, embryos, and somatic cells (Kim et al., 2011, 2010; Lo et al., 2020). Fertilization relies on prototypical Ca^2+^ rises and oscillations, and Zn^2+^ sparks are an egg activation event downstream of this Ca^2+^ release, establishing a functional association between these two divalent cations that continues to grow (Kim et al., 2011). Here, we show that, in addition, these cations actively crosstalk during fertilization and that the fertilization-induced Ca^2+^ oscillations rely on optimized IP_3_R1 function underpinned by ideal Zn^2+^ levels set during oocyte maturation. Future studies should explore if artificial alteration of Zn^2+^ levels can extend the fertile lifespan of eggs, improve developmental competence, or develop methods of non-hormonal contraception.

## Materials and Methods

### Key resources table

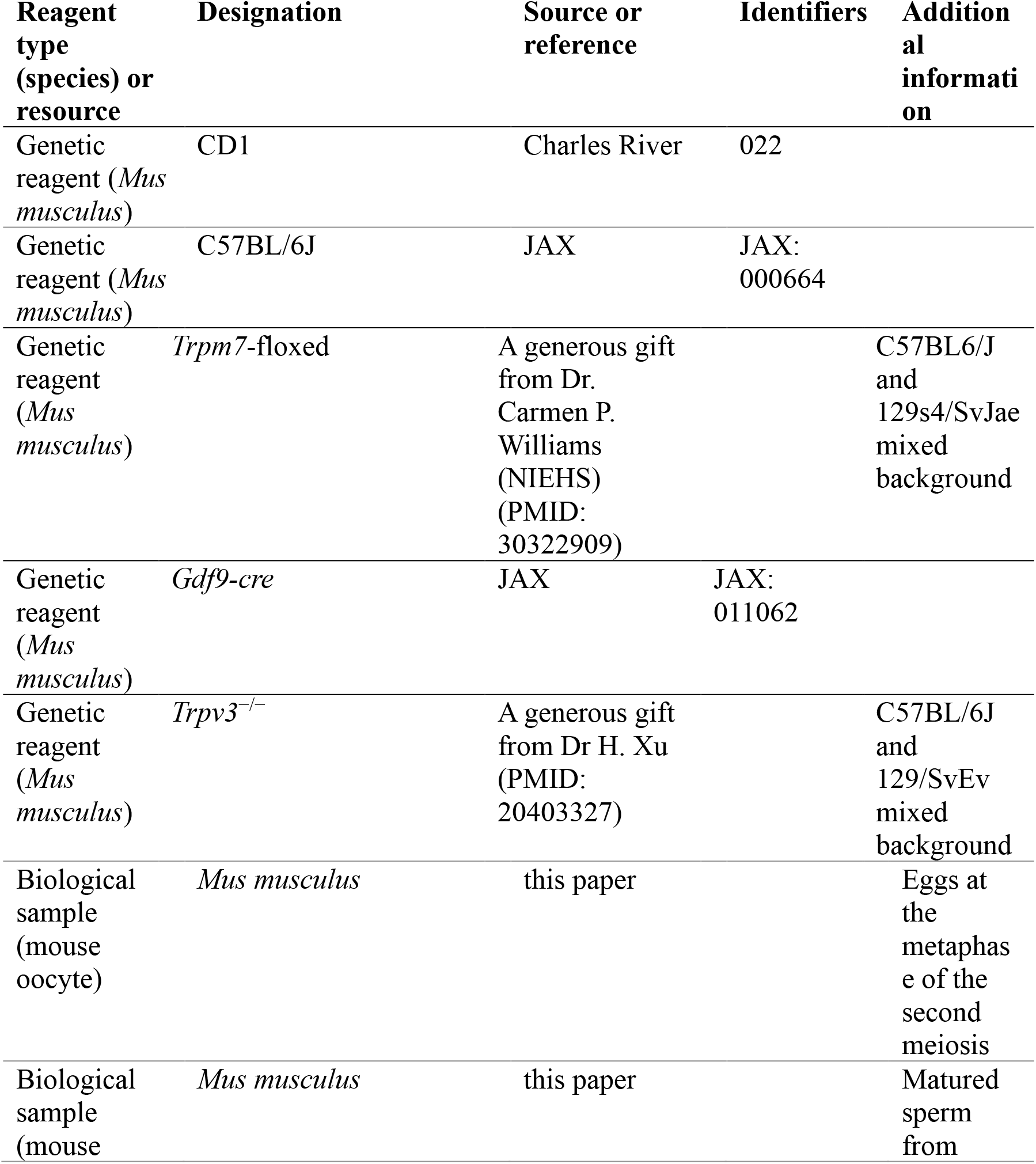

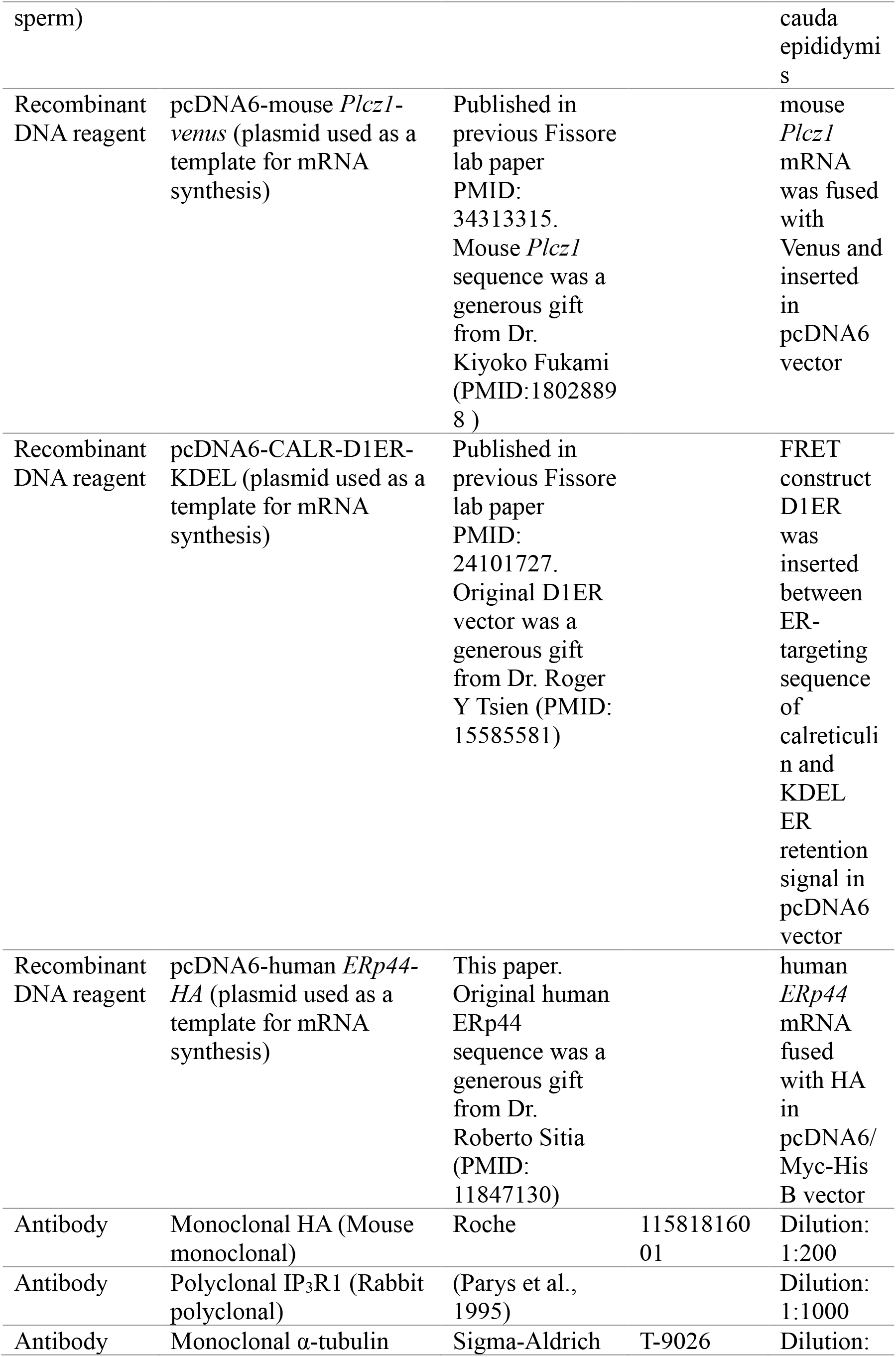

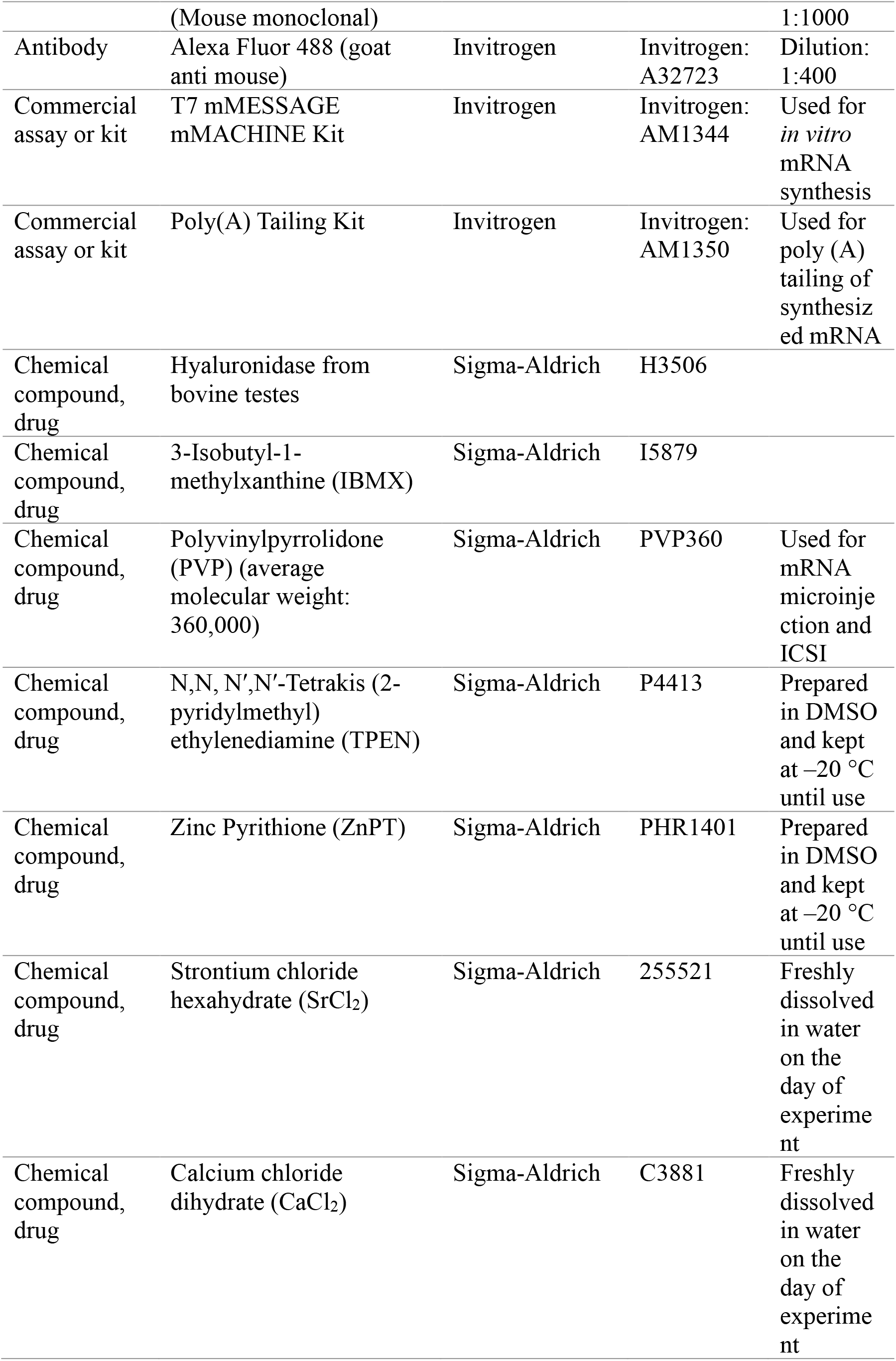

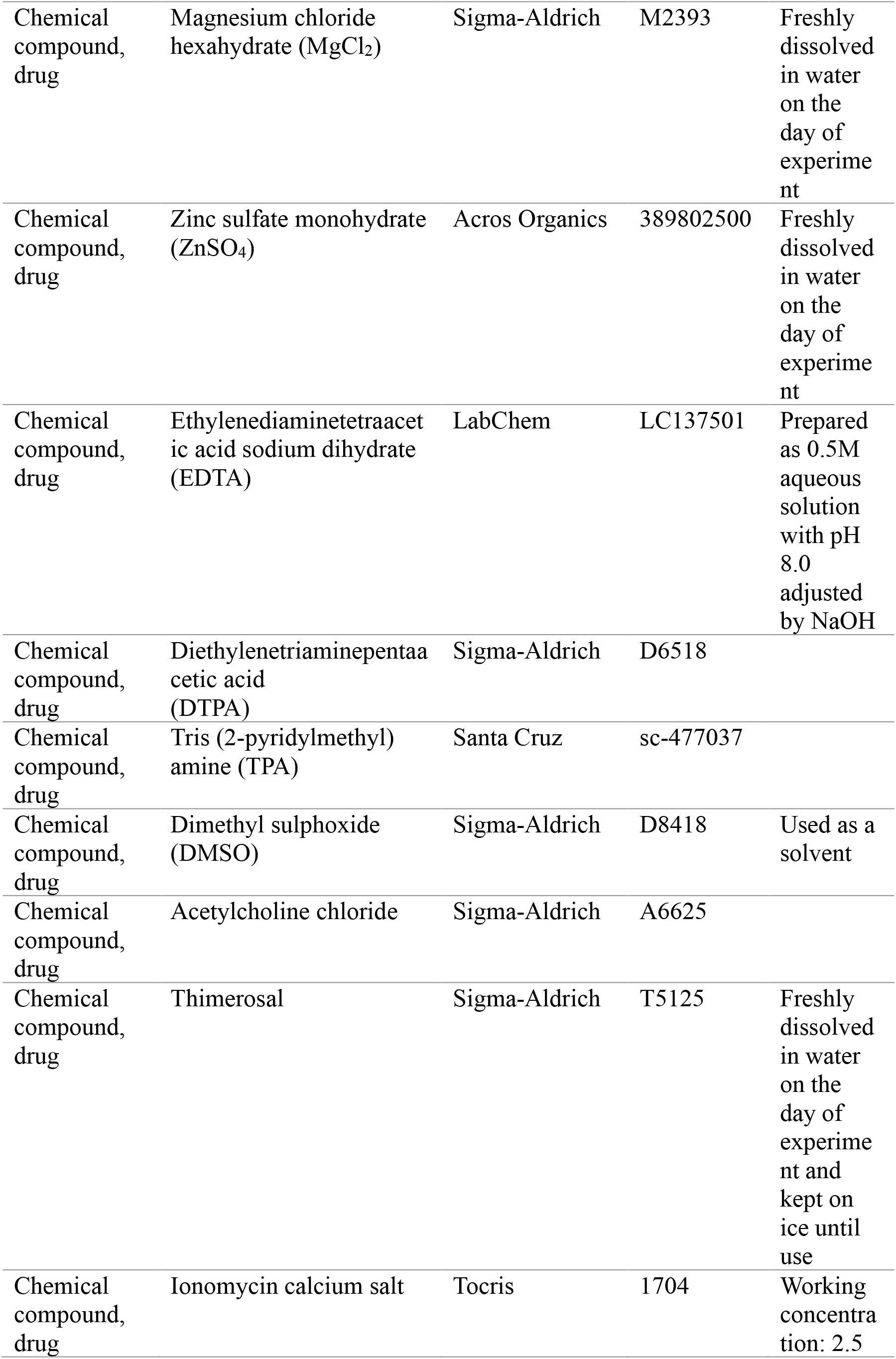

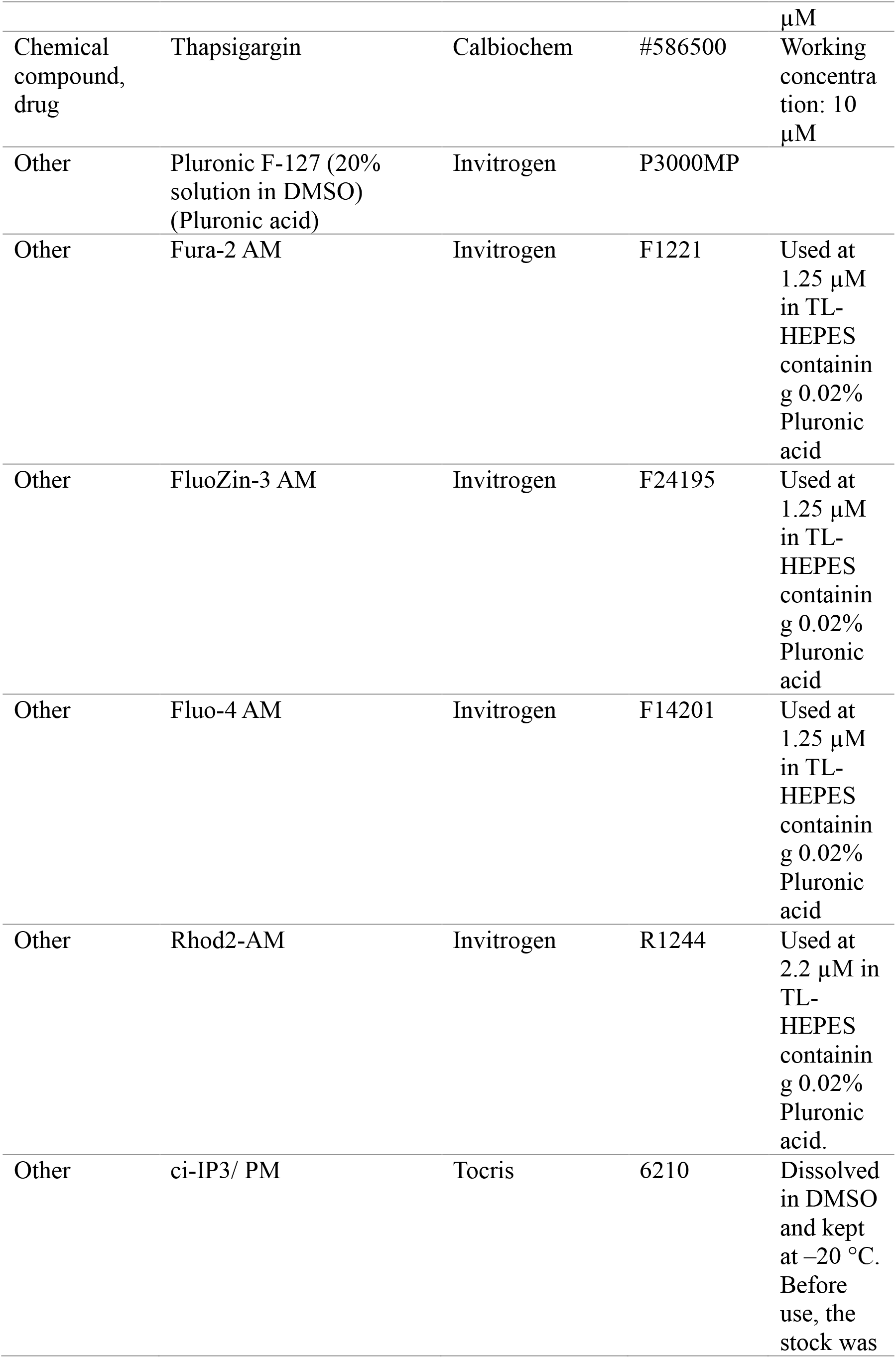

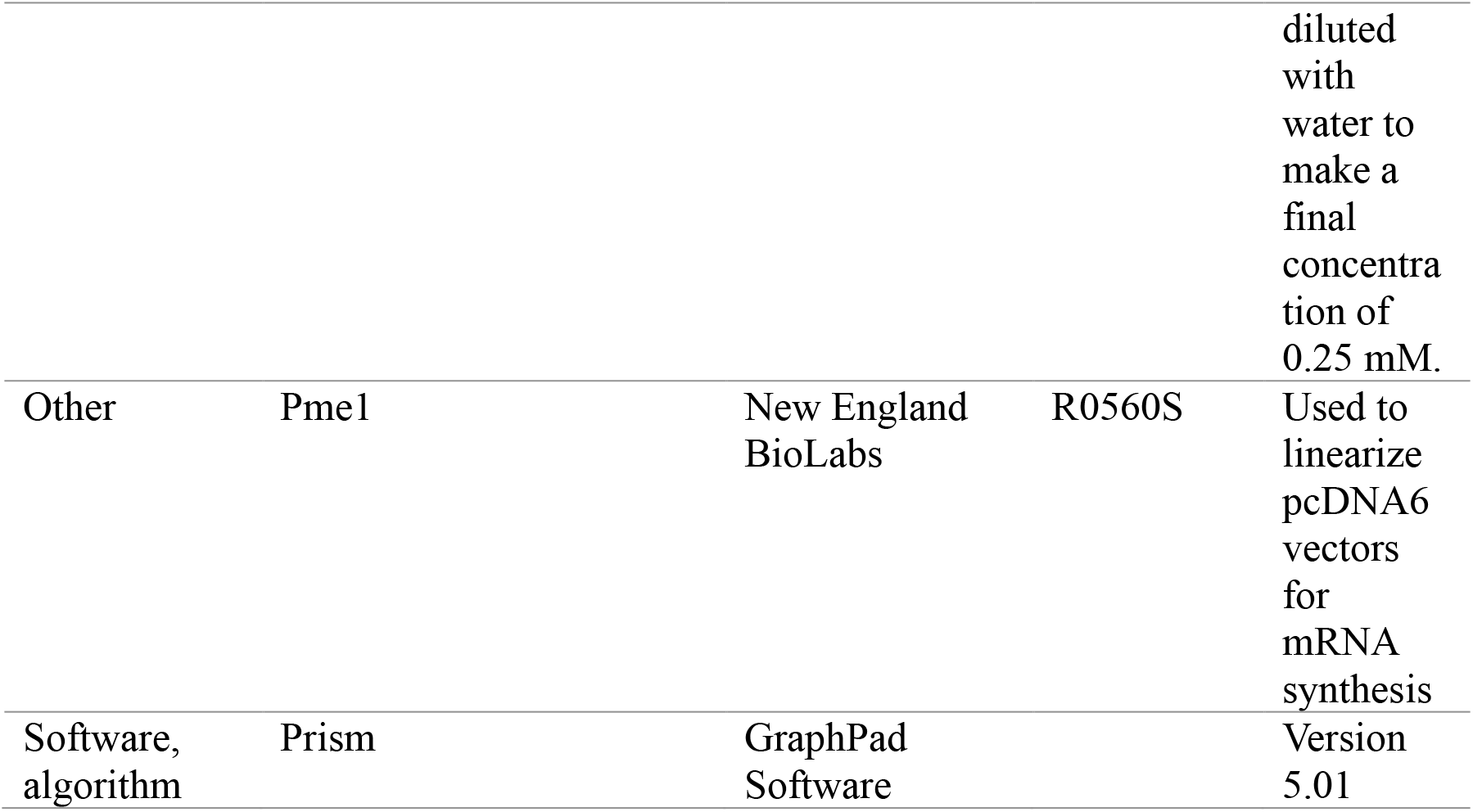

N,N,N′,N′-tetrakis (2-pyridinylmethyl)-1,2-ethylenediamine (TPEN) and Zinc pyrithione (ZnPT) were dissolved in dimethyl sulfoxide (DMSO) at 10 mM and stored at −20°C until use. SrCl_2_, CaCl_2_, ZnSO_4_, and MgCl_2_ were freshly dissolved with double-sterile water at 1M and diluted with the monitoring media just before use. Ethylenediaminetetraacetic acid (EDTA) and diethylenetriaminepentaacetic acid (DTPA) were reconstituted with double-sterile water at 0.5M and 10 mM, respectively, and the pH was adjusted to 8.0. Tris(2-pyridylmethyl) amine (TPA) was diluted in DMSO at 100 mM and stored at −20°C until use. Acetylcholine chloride and Thimerosal were dissolved in double-sterile water at 550 mM and 100 mM, respectively. Acetylcholine was stored at −20°C until use, whereas Thimerosal was made fresh in each experiment.

### Mice

The University of Massachusetts Institutional Animal Care and Use Committee (IACUC) approved all animal experiments and protocols. *Trpm7*-floxed (*Trpm7*^fl/fl^) *Gdf9-Cre* and *Trpv3*^−/−^ mice were bred at our facility. *Trpm7*^fl/fl^ mice were crossed with *Trpv3*^−/−^ to generate *Trpm7*^fl/fl^; *Trpv3*^−/−^ mouse line. Female *Trpm7*^fl/fl^; *Trpv3*^−/−^ mice were crossed with *Trpm7*^fl/fl^; *Trpv3*^−/−^; *Gdf9-cre* male to generate females null for *Trpv3* and with oocyte-specific deletion for *Trpm7*. Ear clips from offspring were collected prior to weaning, and confirmation of genotype was performed after most experiments.

### Egg Collection

All gamete handling procedures are as previously reported by us (Wakai and Fissore, 2019). MII eggs were collected from the ampulla of 6- to 8-week-old female mice. Females were superovulated via intraperitoneal injections of 5 IU pregnant mare serum gonadotropin (PMSG, Sigma, St. Louis, MO) and 5 IU human chorionic gonadotropin (hCG, sigma) at 48hr. interval. Cumulus-oocyte-complexes (COCs) were obtained 13.5 hr. post-hCG injection by tearing the ampulla using forceps and needles in TL-HEPES medium. COCs were treated with 0.26% (w/v) of hyaluronidase at room temperature (RT) for 5 min to remove cumulus cells.

### Intracytoplasmic sperm injection (ICSI)

ICSI was performed as previously reported by us (Kurokawa and Fissore, 2003) using described setup and micromanipulators (Narishige, Japan). Sperm from C57BL/6 or CD1 male mice (7-12 weeks old) were collected from the cauda epididymis in TL-HEPES medium, washed several times, heads separated from tails by sonication (XL2020; Heat Systems Inc., USA) for 5 s at 4°C. The sperm lysate was washed in TL-HEPES and diluted with 12% polyvinylpyrrolidone (PVP, MW = 360 kDa) to a final PVP concentration of 6%. A piezo micropipette-driving unit was used to deliver the sperm into the ooplasm (Primetech, Ibaraki, Japan); a few piezo-pulses were applied to puncture the eggs’ plasma membrane following penetration of the zona pellucida. After ICSI, eggs were either used for Ca^2+^ monitoring or cultured in KSOM to evaluate activation and development at 36.5°C in a humidified atmosphere containing 5% CO_2_.

### Preparation and microinjection of mRNA

pcDNA6-m*Plcζ-mEGFP*, pcDNA6-CALR-D1ER-KDEL, and pcDNA6-*humanERp44-HA* were linearized with the restriction enzyme PmeI and *in vitro* transcribed using the T7 mMESSAGE mMACHINE Kit following procedures previously used in our laboratory (Ardestani et al., 2020). A poly(A) tail was added to the *in vitro* synthesized RNA (mRNA) using Tailing Kit followed by quantification and dilution to 0.5 μg/μL in nuclease-free water and stored at −80°C until use. Before microinjection, m*Plcζ*, D1ER, and *ERp44* mRNA were diluted to 0.01, 1.0, and 0.5 μg/μL, respectively, in nuclease-free water, heated at 95°C for 3 min followed by centrifugation at 13400×*g* for 10 min at 4°C. Cytoplasm injection of mRNA was performed under microscopy equipped with micromanipulators (Narishige, Japan). The zona pellucida and the plasma membrane of MII eggs were penetrated by applying small pulses generated by the piezo micromanipulator (Primetech, Ibaraki, Japan). The preparation of the injection pipette was as for ICSI (Kurokawa and Fissore, 2003), but the diameter of the tip was ∼1 μm.

### Ca_2+_ and Zn_2+_ imaging

Before Ca^2+^ imaging, eggs were incubated in TL-HEPES containing 1.25 μM Fura2-AM, 1.25 μM FluoZin3-AM, or 2.2 μM Rhod2-AM and 0.02% Pluronic acid for 20 min at room temperature and then washed. The fluorescent probe-loaded eggs were allowed to attach to the bottom of the glass dish (Mat-Tek Corp., Ashland, MA). Eggs were monitored simultaneously using an inverted microscope (Nikon, Melville, NY) outfitted for fluorescence measurements. Fura-2 AM, FluoZin3-AM, and Rhod2-AM fluorescence were excited with 340 nm and 380 nm, 480 nm, and 550 nm wavelengths, respectively, every 20 sec, for different intervals according to the experimental design and as previously performed in the laboratory. The illumination was provided by a 75-W Xenon arc lamp and controlled by a filter wheel (Ludl Electronic Products Ltd., Hawthorne, NY). The emitted light above 510 nm was collected by a cooled Photometrics SenSys CCD camera (Roper Scientific, Tucson, AZ). Nikon Element software coordinated the filter wheel and data acquisition. The acquired data were saved and analyzed using Microsoft Excel and GraphPad using Prism software (Ardestani et al., 2020). For Figures 1A, 4A-C, 5A, and 6H-I, values obtained from FluoZin3-AM, Fura2-AM, or Rhod2-AM recordings were divided by the average of the first five recordings for each treatment that was used as the F_0_.

To estimate relative changes in Ca^2+^-ER, emission ratio imaging of the D1ER (YFP/CFP) was performed using a CFP excitation filter, dichroic beamsplitter, CFP and YFP emission filters (Chroma technology, Rockingham, VT; ET436/20X, 89007bs, ET480/40m, and ET535/30m). To measure Ca^2+^-ER and cytosolic Ca^2+^ simultaneously, eggs that had been injected with D1ER were loaded with Rhod-2AM, and CFP, YFP, and Rhod-2 intensities were collected every 20 sec.

### Caged IP_3_

Caged-IP_3_/PM (cIP_3_) was reconstituted in DMSO and stored at −20°C until use. Before injection, cIP_3_ stock was diluted to 0.25 mM with water and microinjected as above. After incubation in KSOM media at 37°C for 1-hr., the injected eggs were loaded with the fluorophore, 1.25 μM Fluo4-AM, and 0.02% Pluronic acid and handled as above for Fura-2 AM. The release of cIP_3_ was accomplished by photolysis using 0.5 to 5-sec pulses at 360 nm wavelengths. Ca^2+^ imaging was as above, but Fluo4 was excited at 488 nm wavelength and emitted light above 510 nm collected as above.

### Western blot analysis

Cell lysates from 20-50 mouse eggs were prepared by adding 2X-Laemmli sample buffer.

Proteins were separated on 5% SDS-PAGE gels and transferred to PVDF membranes (Millipore, Bedford, MA). After blocking with 5% fat-free milk + TBS, membranes were probed with the rabbit polyclonal antibody specific to IP_3_R1 (1:1000; a generous gift from Dr. Jan Parys, Katholieke Universiteit, Leuven, Belgium; Parys et al., 1995). Goat anti-rabbit antibody conjugated to horseradish peroxidase (HRP) was used as a secondary antibody (1:5000; Goat anti-Rabbit IgG (H+L) Cross-Adsorbed Secondary Antibody, HRP; Invitrogen, Waltham, Ma). For detection of chemiluminescence, membranes were developed using ECL Prime (Sigma) and exposed for 1–3 min to maximum sensitivity film (VWR, Radnor, PA). Broad-range pre-stained SDS–PAGE molecular weight markers (Bio-Rad, Hercules, CA) were run in parallel to estimate the molecular weight of the immunoreactive bands. The same membranes were stripped at 50°C for 30 min (62.5 mM Tris, 2% SDS, and 100 mM 2-beta mercaptoethanol) and re-probed with anti-α-tubulin monoclonal antibody (1:1000).

### Immunostaining and confocal microscopy

Immunostaining was performed according to our previous study (Akizawa et al., 2021). After incubation with or without TPEN, MII eggs were fixed with 4% (w/v) paraformaldehyde in house-made phosphate-buffered saline (PBS) for 20 min at room temperature and then permeabilized for 60 min with 0.2% (v/v) Triton X-100 in PBS. Next, the eggs were blocked for 45 min with a blocking buffer containing 0.2% (w/v) skim milk, 2% (v/v) fetal bovine serum, 1% (w/v) bovine serum albumin, 0.1% (v/v) TritonX-100, 0.75% (w/v) glycine in PBS. Eggs were incubated overnight at 4°C with mouse anti-HA antibody (1:200) diluted in blocking buffer. Eggs were washed in blocking buffer 3X for 10 min, followed by incubation at room temperature for 30 min with a secondary antibody, Alexa Fluor 488 goat anti-mouse IgG (H + L) (1:400) diluted in blocking buffer. Fluorescence signals were visualized using a laser-scanning confocal microscope (Nikon A1 Resonant Confocal with six-color TIRF) fitted with a 63×, 1.4 NA oil-immersion objective lens.

### Statistical analysis

Comparisons for statistical significance of experimental values between treatments and experiments were performed in three or more experiments performed on different batches of eggs in most studies. Given the number of eggs needed, WB studies were repeated twice. Prism-GraphPad software was used to perform the statistical comparisons that include unpaired Student’s t-tests, Fisher’s exact test, and One-way ANOVA followed by Tukey’s multiple comparisons, as applicable, and the production of graphs to display the data. All data are presented as mean±s.d. Differences were considered significant at *P* < 0.05.

## Acknowledgments

We thank Ms. Changli He for technical support and Dr. James Chambers for support with confocal microscopy support. We thank all members of the Fissore lab for useful discussions and suggestions. We thank Jan B. Parys, K.U. Leuven, Belgium, for initial discussions and advice.

## Additional information

### Funding Sources

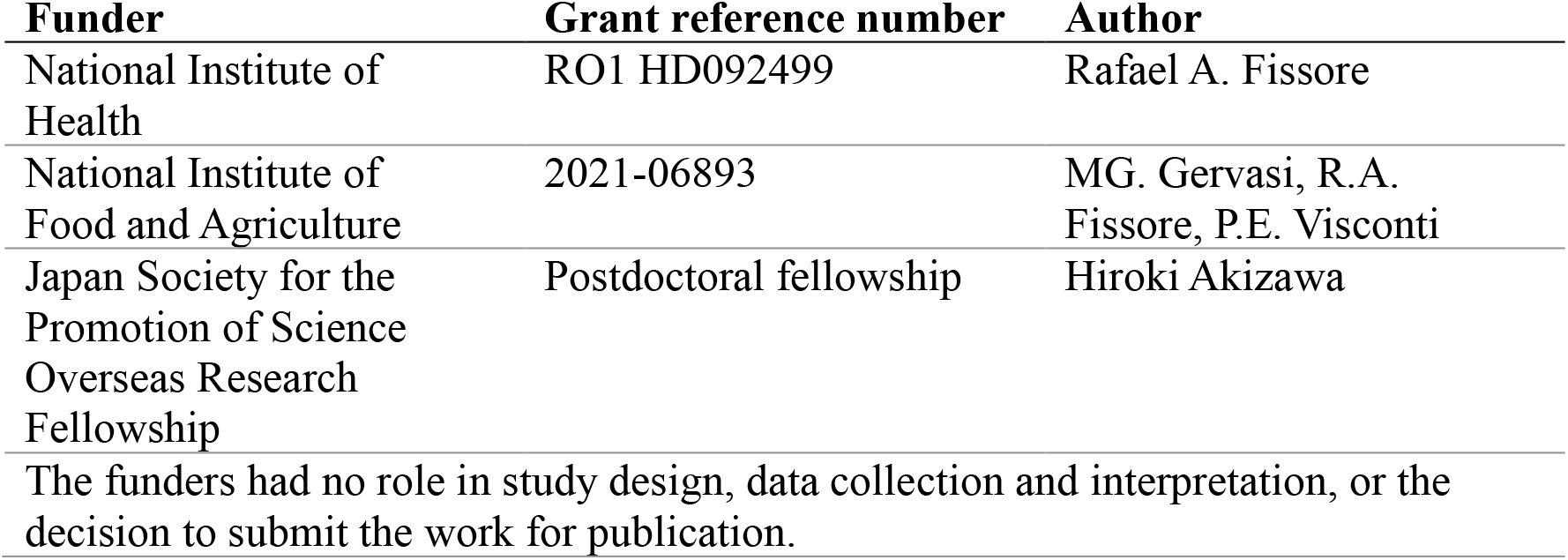

The funders had no role in study design, data collection and interpretation, or the decision to submit the work for publication.

### Author contributions

Hiroki Akizawa, Data curation, Formal analysis, Validation, Investigation, Visualization, Writing—original draft, Writing—review and editing; Emily Lopes, Data curation, Formal analysis, Validation; Rafael A Fissore, Conceptualization, Formal analysis, Supervision, Funding acquisition, Methodology, Writing—original draft, Project administration, Writing—review and editing

---

**Supplementary Figure 1.**
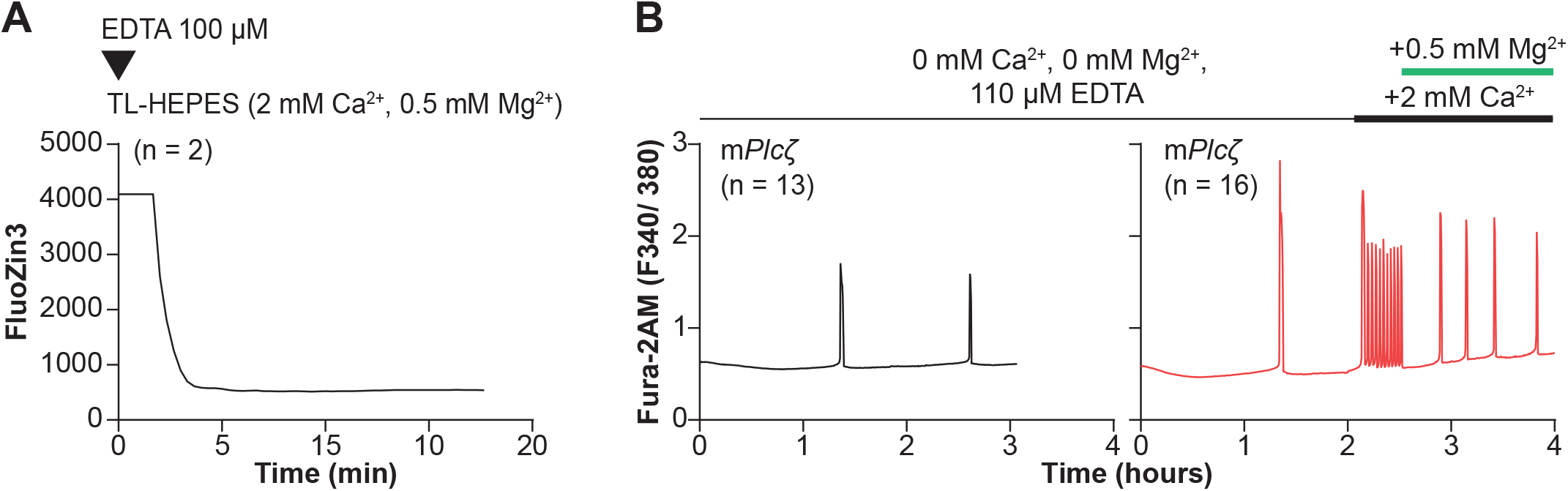
Cell-impermeable chelators effectively reduce Zn_2+_ levels in external media but do prevent initiation or continuation of Ca_2+_ oscillations. (A) A representative trace of FluoZin3 fluorescence in replete monitoring media (TL-HEPES). The media was supplemented with cell-impermeable FluoZin-3, and after initiation of monitoring, the addition of EDTA (100 μM) occurred at the designated point (triangle). (B) The left black trace represents Ca^2^_+_ oscillations initiation by injection of *mPlcζ* mRNA (0.01 μg/μl). The oscillations were monitored in Ca_2+_ and Mg_2+_-free media and in the presence of EDTA (110 μM) to chelate residual divalent cations derived from the water source or reagents used to make the media. The right red trace represents the initiation of oscillations as above, but after a period indicated by the black and green bars, Ca_2+_ and Mg_2+_ were sequentially added back.

**Supplementary Figure 2.**
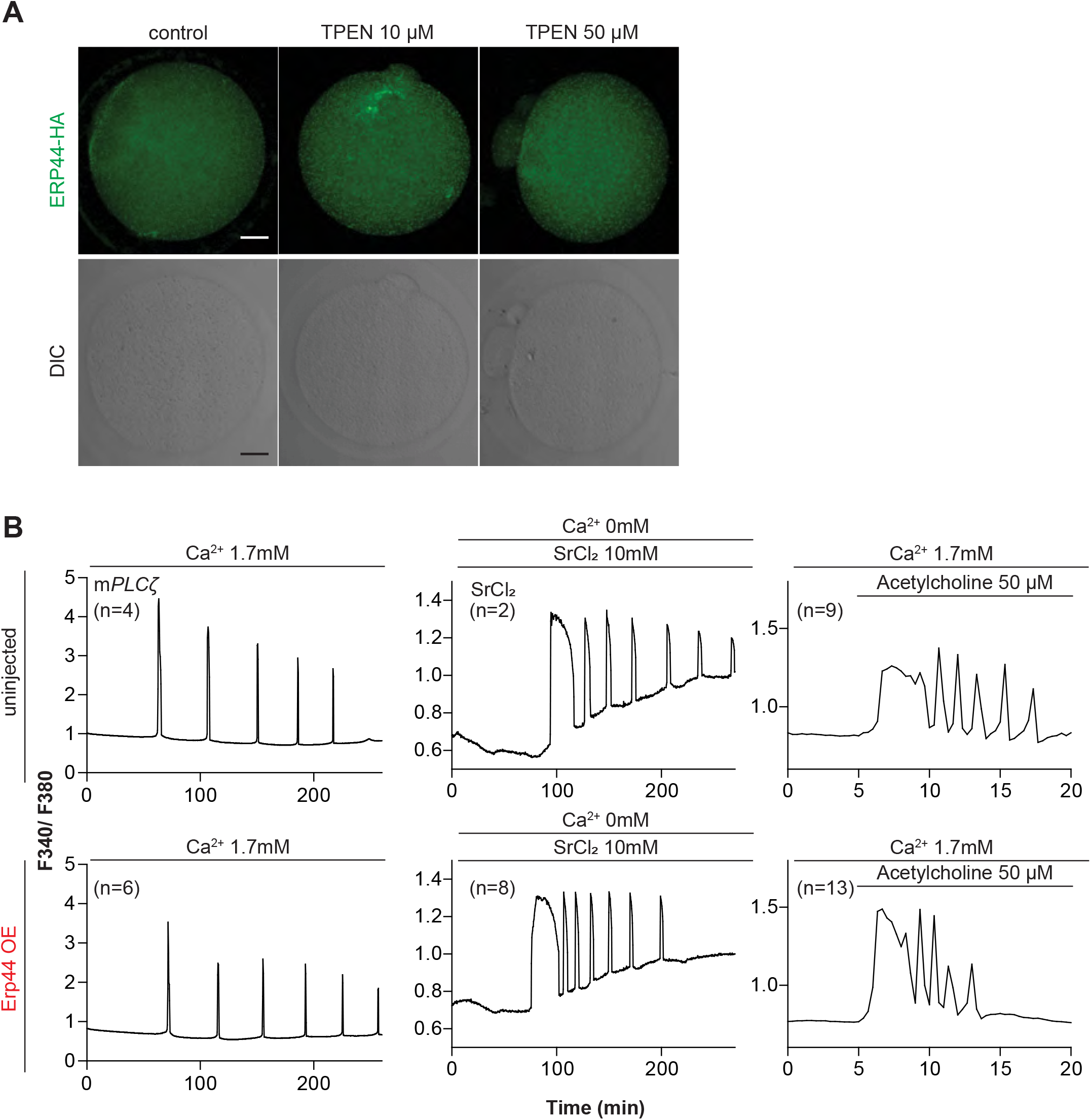
Overexpression of ER accessory protein ERp44 did not change the Ca_2+_ responses initiated by m*Plcζ* mRNA microinjection, Actylcholine, or SrCl_2_. (**A**) Representative immunofluorescent images of MII eggs with overexpression of ERp44. At 5 hr. post microinjection, eggs were treated with 10 or 50 µM of TPEN and incubated for 1 hr, after which they were fixed and stained. An anti-HA antibody was used. Scale bar: 10 µm. (**B**) Representative Ca^2+^ responses induced by m*Plcζ* mRNA microinjection (0.01 µg/ µl-left column), SrCl_2_ (10 mM-center column), and acetylcholine (50 µM-right column) in eggs with (top panels) or without (bottom panels) ERp44 overexpression.

**Supplementary Figure 3.**
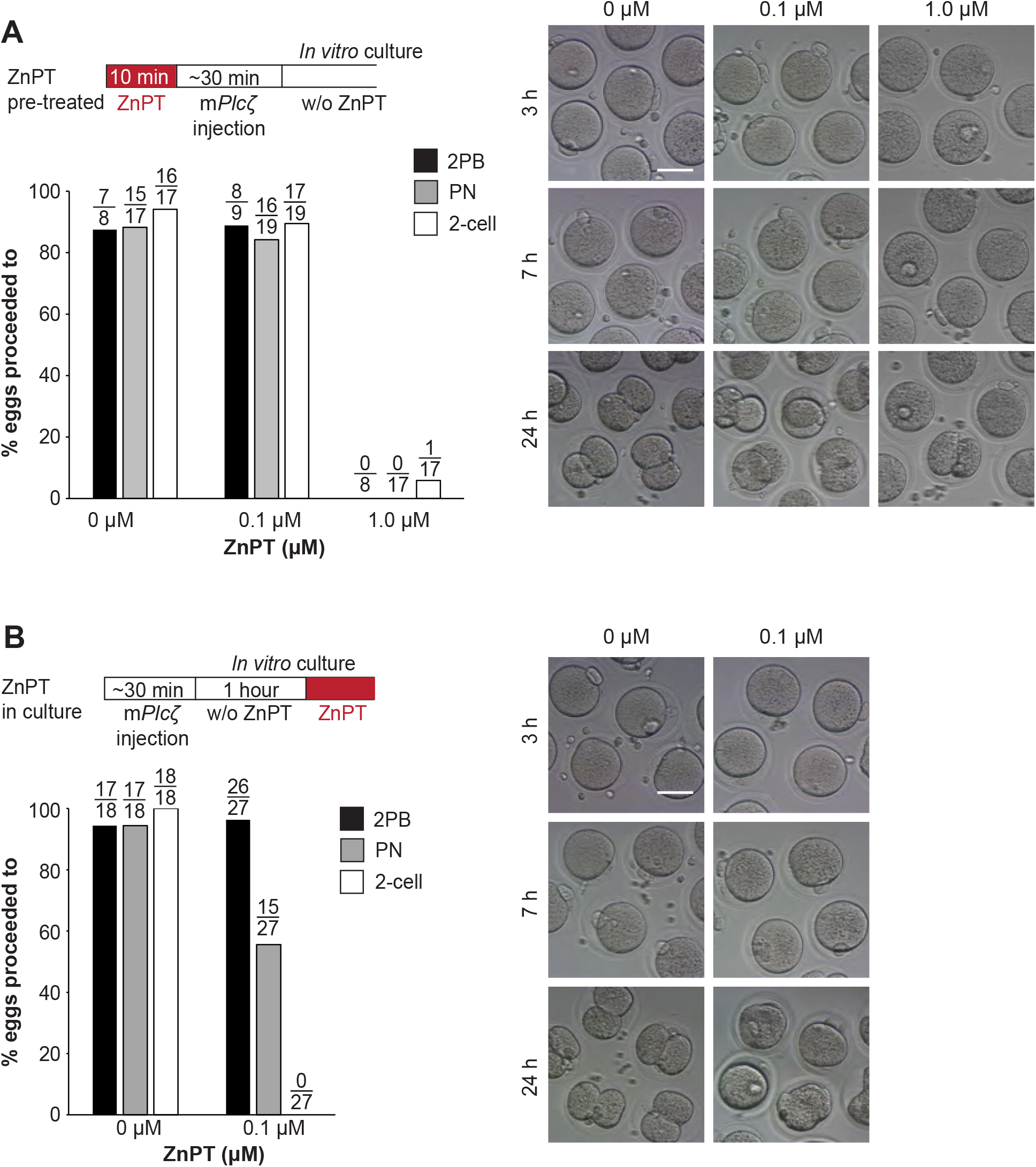
Elevated Zn_2+_ impairs egg activation and the subsequent embryo development. (**A**) MII eggs were incubated in TL-HEPES containing 0, 0.1, or 1.0 µM ZnPT at room temperature for 10 min and washed several times with fresh TL-HEPES and injected with m*Plcζ* mRNA. After it, eggs and zygotes were cultured in KSOM for 24h. PN formation and 2-cell development were checked at 7 and 24h post-microinjection. Bars represent the percentages of injected eggs that reached the PN and the 2-cell stage. Scale bar: 50 μm. (**B**) MII eggs injected with m*Plcζ* mRNA were incubated in KSOM without ZnPT for an hr. and then incubated in KSOM with 0 or 0.1 μM ZnPT for 24h. The second polar body extrusion, PN formation, and 2-cell development were checked at 2.5-, 7- and 24h. post-microinjection. Bars represent the percentages of injected eggs that reached the PN and the 2-cell stage. Scale bar: 50 μm.

